# Circuit decoupling with surrogate compensation for baseline tuning in biophysically detailed neural models

**DOI:** 10.64898/2026.11.13.766494

**Authors:** Nikita Novikov, Scott McElroy, James Chen, Ethan Irby, Salvador Dura-Bernal, Samuel Neymotin

## Abstract

The baseline regime of a neural circuit shapes its dynamical behavior. Establishing an appropriate baseline is therefore an important part of tuning a computational circuit model. This baseline can be specified in terms of target population firing rates and achieved by adjusting the background inputs to each population. Due to recurrent coupling, these inputs need to be tuned jointly, requiring a high-dimensional search that is especially costly for biophysically detailed circuit models. This remains a major bottleneck since current optimization methods are limited in their ability to tackle this problem. Decoupling the circuit by removing all the connections between its cells allows background inputs to be tuned independently, but the solution may not hold when the connectivity is restored. We developed a compensated decoupling method in which missing recurrent inputs are replaced by surrogate inputs from spike generators firing at the corresponding target rates. The resulting input approximates the recurrent drive expected at the target regime, so the background inputs tuned in this setup may remain valid after restoring connectivity. We applied the method to a 43-population biophysically detailed model of the macaque auditory thalamocortical circuit, which had previously proved challenging to fit closely to experimental data. First, we tuned the background input independently for each population in the fully decoupled setup. Next, we tested the solution in the isolated cortical and thalamic subcircuits, preserving recurrent connections within each subcircuit. The cortical subcircuit had two coexisting regimes and settled into the off-target one by default. A gradual transition from surrogate to recurrent self-excitation allowed it to reach the near-target regime. The thalamic subcircuit exhibited a single regime deviating from the target due to imperfect surrogate compensation. Jointly tuning a separate tonic input correction for each thalamic population substantially reduced the errors. Finally, we tested the solution in the full thalamocortical circuit, combining the tuned background inputs, thalamic corrections, and gradual restoration of self-excitation. The full circuit consistently reached a near-target regime across multiple random model realizations. The approach yielded an improved macaque auditory thalamocortical baseline model, providing a community resource for investigating the auditory thalamocortical system’s dynamics. Together, the results demonstrate that our method can facilitate baseline tuning in a biophysically detailed model by enabling decoupled optimization and guiding subsequent refinement.

## Introduction

Biophysically detailed circuit models bring together data on neuronal properties, synaptic interactions, and circuit anatomy to support in silico experiments (Arkhipov et al., 2018; Billeh et al., 2020; Dura-Bernal et al., 2023a; Isbister et al., 2026). Direct measurements constrain some model parameters, while others should be refined by fitting simulated activity to physiological recordings. Such tuning often requires a high-dimensional search, with candidate parameter combinations evaluated through computationally expensive simulations. Several optimization approaches have been applied to biophysically detailed circuit models (Dura-Bernal et al., 2019, 2023a; Ranak and Anderson, 2026), and new methods continue to emerge (Deistler et al., 2025). Nevertheless, reproducing physiological circuit activity remains difficult and requires tuning procedures adapted to the model and available data (Dura-Bernal et al., 2024; Tolley et al., 2024).

Establishing a physiologically appropriate baseline regime in a circuit model is important for producing realistic responses to incoming stimuli. Experimental studies have demonstrated the dependence of these responses on the baseline regime at both the single-neuron (Chance et al., 2002; Fellous et al., 2003) and the population level (Petersen et al., 2003; Curto et al., 2009; Zagha et al., 2013; McGinley et al., 2015; Gutnisky et al., 2017). Nonlinear recurrent network models likewise show that changes in the baseline regime can alter effective coupling between cells even when synaptic weights remain fixed (Ahmadian et al., 2013; Rubin et al., 2015; Hennequin et al., 2018).

Baseline activity can be set by tuning either recurrent synaptic weights (Arkhipov et al., 2018; Billeh et al., 2020; Dura-Bernal et al., 2023a) or population-specific background inputs that represent aggregate synaptic activity from sources outside the modeled circuit (Jiang et al., 2024; Isbister et al., 2026). Weight tuning can yield more self-sufficient models, but typically involves many adjustable parameters and requires computationally demanding optimization. Moreover, fitting the baseline through this approach alone may alter circuit interactions in ways inconsistent with experimental evidence. By comparison, background-input tuning involves fewer parameters and allows the baseline to be set without changing the weights, although the biological sources of such inputs are often only partly specified. Each tuning strategy therefore has its own strengths and limitations, and the choice depends on the available constraints and modeling objectives. In this work, we develop a tuning framework within the background-input approach.

Despite its smaller parameter set, background-input tuning remains a multidimensional problem. Changing the input to one population affects the whole circuit through recurrent connectivity, so population-specific inputs have to be tuned jointly. One way to simplify this problem is to decouple the circuit by removing its recurrent connections and tune each population’s background inputs independently. However, due to omitting the recurrent inputs that cells normally receive from each other, the background inputs tuned in the decoupled setup may yield a different regime when recurrent connectivity is restored. A possible solution was proposed by Isbister et al. (2026): the authors first measured how each population’s activity depended on its background input in the decoupled setup, and then iteratively estimated the relationship between the activity produced in the decoupled and recurrent circuits. Together, these relationships allowed them to select background inputs expected to produce the required baseline with restored recurrent connectivity.

Here we present an alternative method of background-input tuning in multi-population models, based on circuit decoupling with surrogate compensation for the omitted recurrent connections. The method allows each population’s background input to be tuned independently under synaptic conditions approximating those expected in the recurrent circuit operating in the target baseline regime. We apply the method to a modified version of our previously published 43-population model of the macaque auditory thalamocortical circuit (Dura-Bernal et al., 2023a) and assess whether independently tuned background inputs can support the chosen baseline regime in the recurrently connected circuit. We also examine how surrogate-compensated decoupling can help identify and resolve issues encountered in the tuning process.

## Results

### Circuit decoupling with surrogate compensation

We developed a method for tuning background inputs in multi-population models based on circuit decoupling with automatic replacement of removed recurrent connections by compensatory surrogate inputs. In a fully decoupled circuit, cells do not affect each other through recurrent connections, so the background inputs can be tuned independently for each population. This transforms a high-dimensional optimization problem into a set of independent, low-dimensional problems that can be solved in parallel. We constructed the surrogate inputs to approximate the synaptic inputs that cells would receive in the original circuit if its populations fired at their target rates. Our rationale was that background inputs tuned in a decoupled model with such surrogate compensation could provide candidate solutions for the original model as well, bringing the original recurrently connected circuit close to the target regime.

The method is illustrated in **Fig. 1** using a hypothetical circuit containing two populations, *P*_1_ and *P*_2_. The original circuit contains within- and cross-population recurrent projections *P*_*m*_ → *P*_*n*_ (*m, n* = 1,2), each comprising a set *C*_*nm*_ of individual cell-to-cell connections (**Fig. 1A**). In addition, each cell receives two background synaptic inputs, one excitatory and one inhibitory, driven by independent Poisson spike generators. The background input rates and synaptic weights are defined separately for each population and shared among its cells. We keep the synaptic weights fixed and adjust the background rates to make *P*_1_ and *P*_2_ fire at their respective target mean rates, 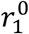 and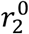.

**Figure 1.**
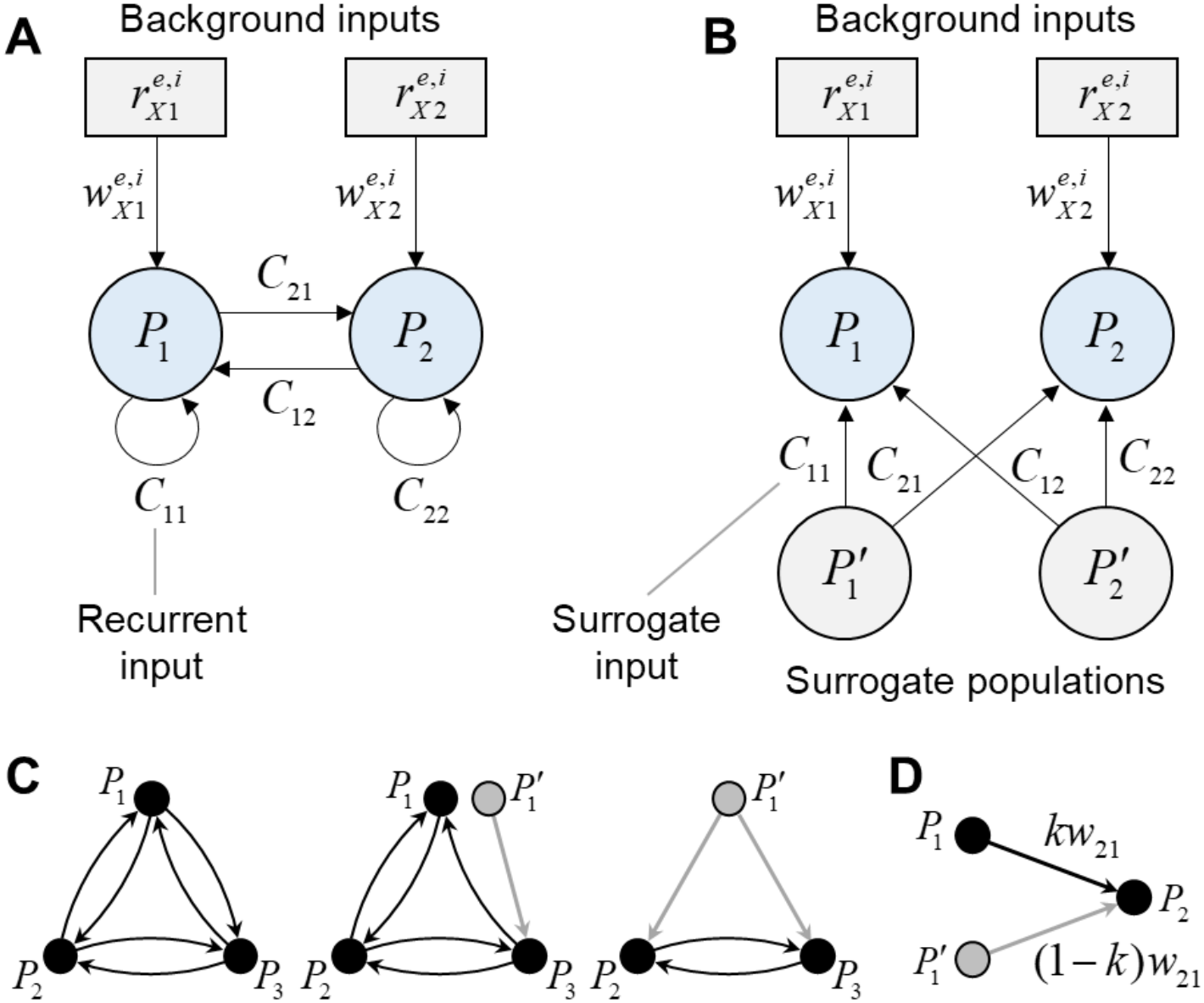
Circuit decoupling with surrogate compensation. **(A)** Original circuit with two populations *P*_1_ and *P*_2_, coupled through recurrent projections *C*_*nm*_ (*m, n* = 1,2). Each population receives background excitatory and inhibitory Poisson inputs, parameterized by firing rates 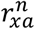 and synaptic weights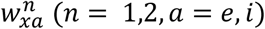 . **(B)** Decoupled open-loop version of the same circuit. Recurrent inputs are replaced by inputs from surrogate populations 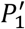 and 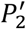 composed of independent Poisson spike generators. The surrogate inputs have the same connectivity properties *C*_*nm*_ (*m, n* = 1,2) as in the original circuit. This transformation decouples the optimization problem, allowing the background input parameters to be tuned independently for each population. **(C)** Partially surrogate setup. Left – original circuit; middle – circuit with frozen *P*_1_ → *P*_3_ projection (replaced by a surrogate); right – isolated *P*_2_ + *P*_3_ subcircuit with *P*_1_ → *P*_2_ and *P*_1_ → *P*_3_ projections replaced by surrogates. **(D)** Semi-surrogate setup. Weight of *P*_1_ → *P*_2_ projection is reduced from *w*_21_ to *kw*_21_ (0 < *k* < 1), and the missing part is compensated by a surrogate 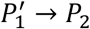 input with weight (1 − *k*)*w*_21_.

A fully decoupled version of the circuit is shown in **Fig. 1B**. To compensate for the omitted recurrent inputs, we introduce a surrogate population 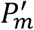 for each original population *P_m_*, consisting of independent Poisson spike generators that fire at the target rate 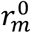 assigned to *P_m_*. We then replace every recurrent projection *P*_*m*_ → *P*_*n*_ with a corresponding surrogate input 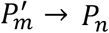 . We connect the surrogate units of 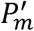 to cells of *P*_*n*_ following the same rules as for the original *P*_*m*_ → *P*_*n*_ projection, preserving connection probabilities and the distributions of synaptic weights and transmission delays. We position the surrogate units according to the same spatial distribution as the cells of *P*_*m*_, thus preserving distance-dependent terms in the connectivity rules. In the resulting circuit, the original cells and their background inputs remain unchanged, while their recurrent inputs are replaced by surrogate inputs from spike generators.

Surrogate spike trains do not depend on the activity of *P*_1_ and *P*_2_, so changing either population’s background inputs in the decoupled circuit does not affect the other population. Therefore, the background inputs to *P*_1_ and *P*_2_ can be tuned separately. If both populations fired at their target rates in the original circuit, the recurrent projections in **Fig. 1A** and the corresponding surrogate projections in **Fig. 1B** would have the same mean presynaptic firing rates. This equality suggests that background inputs tuned in the decoupled setup may also produce a near-target regime in the original circuit. For a circuit with *N* populations, this approach would allow replacing a 2*N*-dimensional search of background-input parameters with *N* independent two-parameter searches. However, whether the resulting background rates actually yield a near-target regime in the recurrently connected circuit remains to be tested.

Our framework also enables more detailed circuit investigation through partial decoupling, in which selected recurrent projections are replaced by surrogate inputs while the remaining projections are preserved. This provides two complementary ways to examine circuit dynamics. First, we can assess the contribution of individual projections or groups of projections by replacing them with surrogate inputs while leaving the rest of the recurrent circuit intact. Second, we can isolate a selected subcircuit by retaining its internal recurrent projections and replacing inputs from outside the subcircuit with surrogates. **Figure 1C** illustrates both operations for a three-population circuit.

Finally, our framework supports semi-surrogate projections, which combine recurrent and surrogate inputs (**Fig. 1D**). For each selected projection, we retain the recurrent connections and independently generate a surrogate counterpart using the same connectivity rules. We scale the synaptic weights of the recurrent and surrogate projections by *k* and (1 − *k*), respectively, where 0 ≤ *k* ≤ 1. Thus, *k* = 0 corresponds to fully surrogate input, while *k* = 1 corresponds to the original recurrent projection. By gradually increasing *k* from 0 to 1, we can examine how the circuit responds as recurrent feedback is restored and whether a near-target regime can be maintained.

### Background input tuning in a fully decoupled circuit

We applied our method to a modified version of our previously published biophysical model of the macaque auditory thalamocortical circuit, containing 43 recurrently connected populations (Dura-Bernal et al., 2023a). Our objective was to establish a baseline regime in which each population’s mean firing rate matched a chosen target. The target firing rates for all populations are listed in **Table S1**.

The version used here differed from the published model in its background-input configuration: each neuron received excitatory and inhibitory Poisson inputs representing synaptic activity from sources outside the modeled circuit. Neurons within a population shared the two input rates but received independently generated spike trains. We held both recurrent and background-input synaptic weights fixed and tuned only the two Poisson rates for each population. Across the 43 populations, this gave 86 tunable background rates.

We first searched for background rates that would make the populations fire at their targets in a fully decoupled version of the circuit. Each modeled population retained its background inputs, while its recurrent inputs were replaced by inputs from surrogate populations. The spike generators in each surrogate population fired at the target mean rate assigned to the corresponding modeled population. **Figure 2A** illustrates this configuration for one population, *P* = IT5A, with excitatory and inhibitory background rates 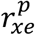 and 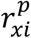, respectively, grouped as 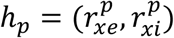. A1^′^ represents the surrogate copy of the entire circuit that supplies compensating inputs to *P*. Because the surrogate inputs were independent of the modeled populations’ activity, adjusting one population’s background rates left the inputs to other populations unchanged. The task of choosing 86 background rates could therefore be carried out as 43 independent 2-D searches, each over the excitatory and inhibitory background rates of one population.

**Figure 2.**
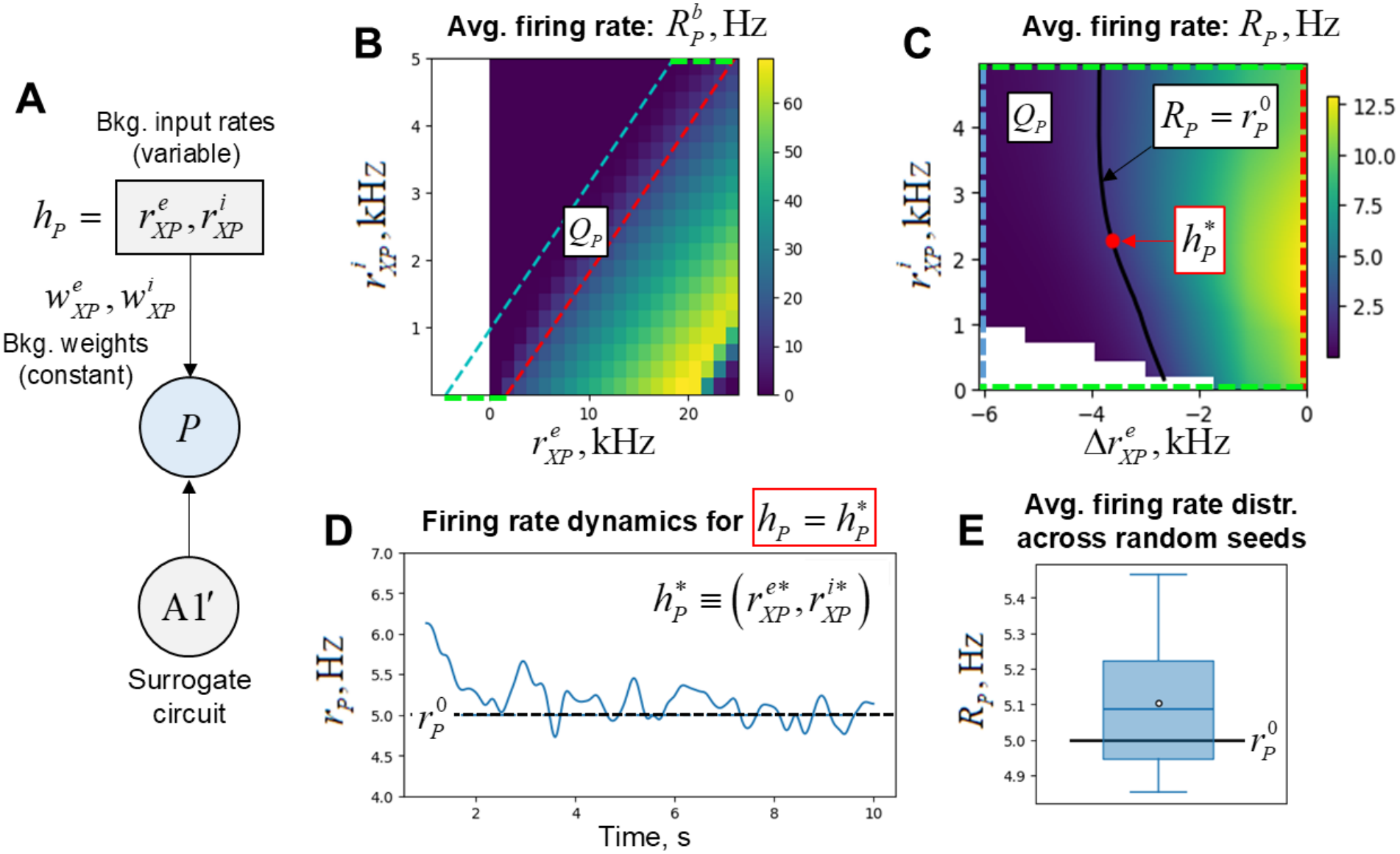
Population-specific search for background input rates in the fully decoupled circuit. The procedure is illustrated for one population *P* = IT5A. **(A)** Schematic of the tuning setup. Population *P* receives excitatory and inhibitory background Poisson inputs 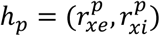 through fixed synaptic weights 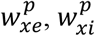 . All recurrent inputs to *P* are replaced by inputs from a surrogate copy of the full circuit (A1^′^), composed of Poisson spike generators firing at the corresponding population target rates. **(B)** Coarse-scan map 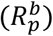 showing the mean firing rate of *P* measured for each combination of background rates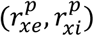 . The parallelogram *Q*_*p*_ encompasses the region where 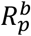 is close to the target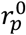. **(C)** Fine-scan map *R*_*p*_ showing the mean firing rates measured within *Q*_*p*_, shown in transformed coordinates in which *Q*_*p*_ becomes a rectangle. Black contour denotes the set of all possible inputs that yield the target firing rate: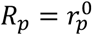 . We chose the final background solution 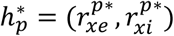 as the midpoint of this contour. **(D)** Firing rate dynamics of *P* obtained with the chosen background input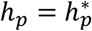, showing convergence toward the target rate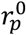 . **(E)** Distribution of the mean firing rates of *P* across 15 trials with random model realizations and the same input 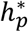.

We performed each of these searches in two stages: a coarse scan followed by a finer scan. **Figure 2B,C** shows the resulting firing rate maps for the illustrated population *P*. For the coarse scan, we simulated the fully decoupled circuit with background rates sampled from a grid spanning a broad rectangular region of the 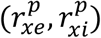 plane and measured the population’s mean firing rate for each combination. The coarse map showed a narrow, approximately diagonal band of background rates for which the population fired near its target, 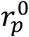 (**Fig. 2B**). We defined a parallelogram *Q*_*p*_ around a portion of this band and performed a finer scan within it, using transformed coordinates in which *Q*_*p*_ appeared as a rectangle (**Fig. 2C**). From the finer measurements, we identified a contour along which the population’s mean firing rate equaled its target. We selected the background-rate pair at the midpoint of this contour within *Q*_*p*_ as the solution and denote this pair by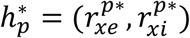.

The searches yielded a set of selected background rates for all 43 populations, which we denote by *H*^*\**^ (**Table S3**). To test these rates across different model realizations, we ran 15 fully decoupled circuits, holding *H*^*\**^ fixed while resampling the connectivity and Poisson spike trains. For the illustrated population *P* = IT5A, the firing rate fluctuated around its target after an initial transient (**Fig. 2D**). Its mean firing rate remained near the target across the 15 trials (**Fig. 2E**). For each population and trial, we calculated firing rate error as the difference between the simulated mean firing rate and the corresponding target rate. The resulting error distributions across trials had reasonably small means and standard deviations for all populations (**Tables S5 and S6**, All-surr. columns).

After tuning the background inputs in the fully decoupled setup, we proceeded to test whether the tuned inputs would also produce near-target firing rates when recurrent connections were restored. We began by testing partially decoupled models in which either the cortical or thalamic subcircuit retained its internal recurrent connections, while inputs from the rest of the network were replaced by surrogates.

### Cortex

#### Fully connected circuit

We began evaluation of the obtained background-input solution starting from the isolated cortical subcircuit. We compared simulation results obtained under the same background input in the fully decoupled cortical subcircuit (**Fig. 3A**) and in the recurrently connected cortical subcircuit with surrogate thalamocortical inputs (**Fig. 3B**). For each configuration, we ran 15 simulation trials with different random model realizations. For each trial, we calculated the mean firing rates of the cortical populations and their deviations from the target rates.

**Figure 3.**
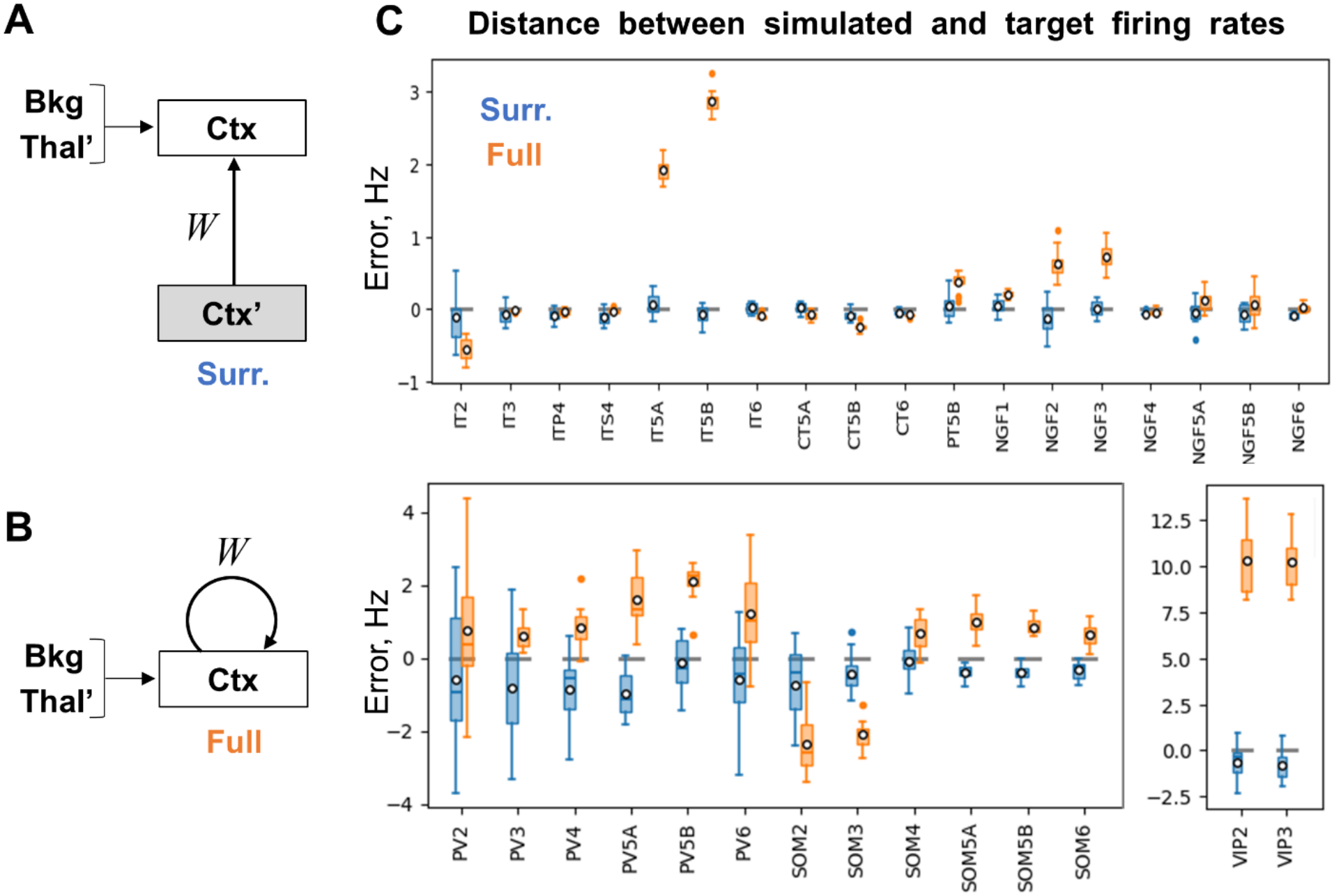
Firing rate errors in the fully decoupled and recurrently connected cortical subcircuits. **(A)** Schematic of the fully decoupled cortical subcircuit. Ctx – original cortical populations; Ctx’ and Thal’ – surrogate copies of cortical/thalamic populations comprised of Poisson spike generators firing at the target rates; Bkg – background input. All non-background inputs of Ctx cells are surrogate, i.e., originate from Ctx’/Thal’ units. **(B)** Schematic of the recurrently connected cortical subcircuit with surrogate thalamic input. **(C)** Distributions of the firing rate errors across 15 random model realizations, calculated as the difference between the simulated mean firing rate of each population and its target rate. Blue – fully decoupled cortical subcircuit shown in A, orange – recurrently connected subcircuit shown in B. Boxes – interquartile ranges (IQR), horizontal lines – medians, white circles – means, whiskers – most extreme samples within 1.5 IQR of the box edges, colored dots – outliers.

The resulting error distributions across trials for the two model configurations are shown in **Figure 3C**. In the fully decoupled circuit, the errors remained close to zero (blue bars), as expected for a network setup identical to the one used for the background tuning step. In the recurrently connected circuit, the errors were substantially larger (orange bars; **Table S5**, Original vs. All-surr.). Particularly strong deviations from the target were observed in IT5A, IT5B, VIP2, and VIP3 populations.

There are several possible explanations for the observed errors. First, the near-target regime could have disappeared or become unstable when recurrent connections were restored, causing the circuit to settle into a different state. Second, the regime could have remained stable but shifted away from the target rates. Third, a stable near-target regime could have coexisted with the observed distant regime, with the state reached depending on the initial conditions. We performed additional investigation to identify the main cause of the observed errors.

#### Connected circuit with frozen self-excitation

We first examined the role of recurrent excitatory-to-excitatory (E-E) connections, since excitatory feedback is often associated with loss of stability or emergence of multiple stable regimes. To assess their contribution, we considered a cortical circuit configuration with all E-E connections replaced by surrogate inputs (“frozen”) and the remaining intracortical connections kept recurrent (**Fig. 4A**).

**Figure 4.**
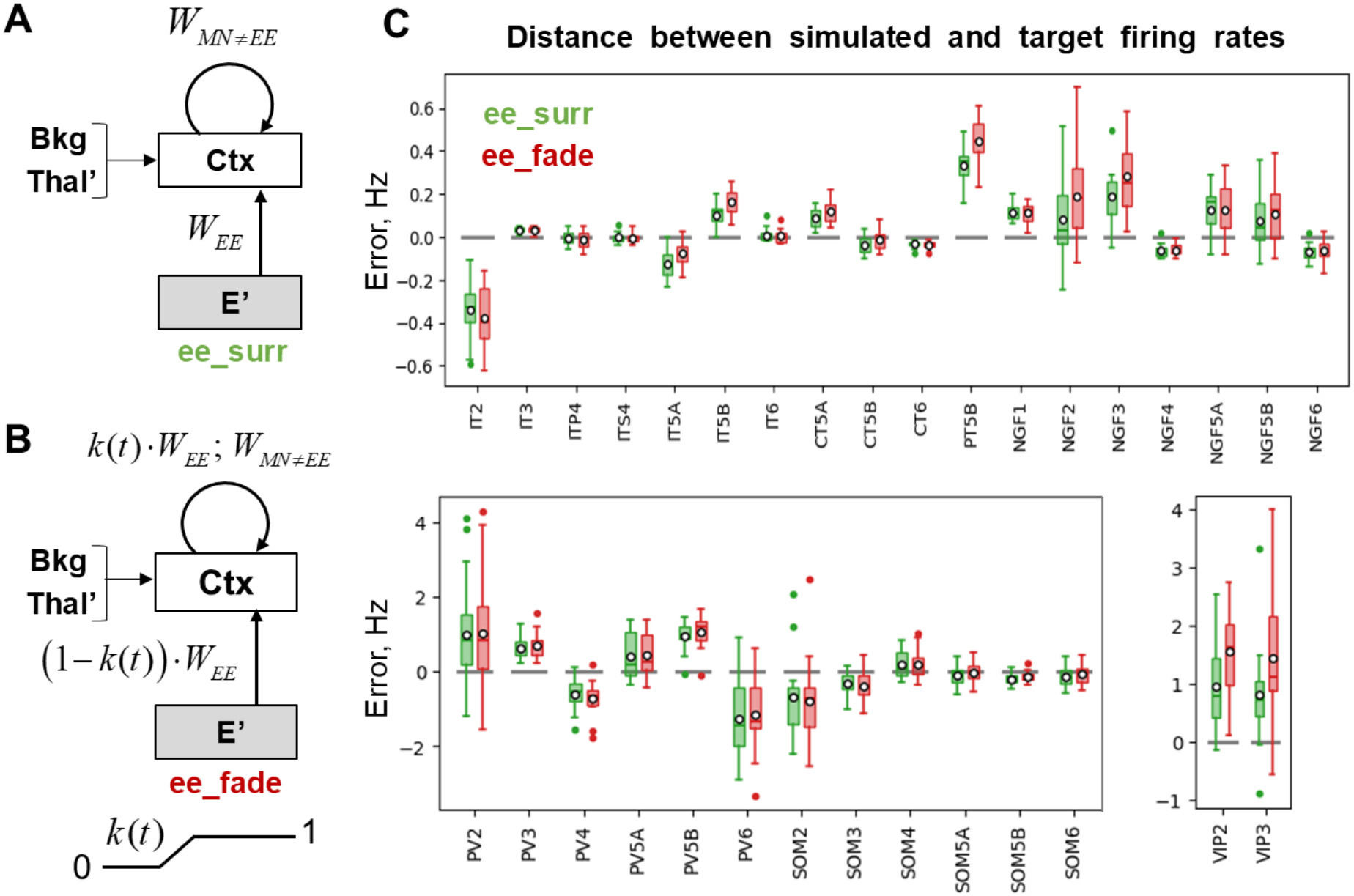
Gradual restoration of recurrent excitatory-to-excitatory connections in the cortical subcircuit. **(A)** Schematic of the cortical subcircuit with frozen excitatory-to-excitatory (E-E) projections. Ctx, Thal^′^, and Bkg are defined as in **Fig. 3**; E^′^ denotes surrogate copies of excitatory cortical populations. E-E projections (*W*_*EE*_) are replaced by surrogate inputs from E^′^, while all other intracortical projections (*W*_*MN*_, *MN* ≠ *EE*) remain recurrent. **(B)** Schematic of the fading procedure, in which surrogate E-E inputs are gradually replaced by recurrent inputs. The synaptic weights of the recurrent and surrogate E–E projections are scaled by *k*(*t*) and 1 − *k*(*t*), respectively. As *k*(*t*) increases from 0 to 1, the circuit transitions from the configuration shown in panel A to the recurrently connected configuration shown in **Fig. 3B. (C)** Distributions of firing rate errors across 15 random model realizations. Green – cortical subcircuit with frozen E-E projections (panel A); red – cortical subcircuit after gradual restoration of recurrent E-E projections (panel B). Boxplot conventions are the same as in **Fig. 3C**.

Across 15 trials, the firing rate errors remained relatively small (**Fig. 4C**, green bars), in contrast to the substantially larger errors observed in the fully interconnected circuit (**Fig. 3C**, orange bars; **Table S5**, Original vs. E-E surr.). These results indicate that recurrent E-E connections played a major role in the observed deviations from the target rates.

#### Fading from frozen to recurrent self-excitation

We next investigated whether the near-target regime obtained with frozen E-E connections could be maintained after these connections were restored. We first ran the circuit with frozen E-E connections for 3 seconds, allowing its activity to settle near the target rates. Over the following 2 seconds, we faded out the surrogate E-E inputs while fading in the corresponding recurrent connections (**Fig. 4B**). We then continued the simulation with all intracortical connections recurrent and calculated the resulting firing rate errors.

Across 15 trials, the errors remained small after the transition to the fully recurrent configuration (**Fig. 4C**, red bars), and the resulting error distributions were broadly similar to those obtained with frozen E-E connections (green bars). Importantly, the final circuit configuration was the same as in the trials with all intracortical connections being recurrent from the simulation outset, but the error distributions differed substantially (red bars in **Fig. 4C** vs. orange bars in **Fig. 3C**; **Table S5**, E-E faded vs. Original). These results indicate that the fully recurrent circuit could sustain both near-target and distant regimes, depending on its preceding dynamics. Gradual restoration of E-E connections allowed the circuit to successfully reach the near-target regime, with the mean errors within ±2 Hz for all cortical populations.

### Thalamus

#### Fully connected circuit

Having evaluated the background solution in the cortical subcircuit, we next turned to the complementary case: the thalamic subcircuit with surrogate cortical inputs. We compared simulation results obtained in the fully decoupled (**Fig. 5A**) and recurrently connected (**Fig. 5B**) thalamic subcircuits. As in the cortical analysis, we ran 15 simulation trials with different random model realizations for each configuration. The resulting error distributions are shown in **Fig. 5D**. In the fully decoupled circuit, the errors remained close to zero (blue bars). In the recurrently connected circuit, the errors were substantially larger (orange bars), particularly in the TC and HTC populations (**Table S6**, Original vs. All-surr.).

**Figure 5.**
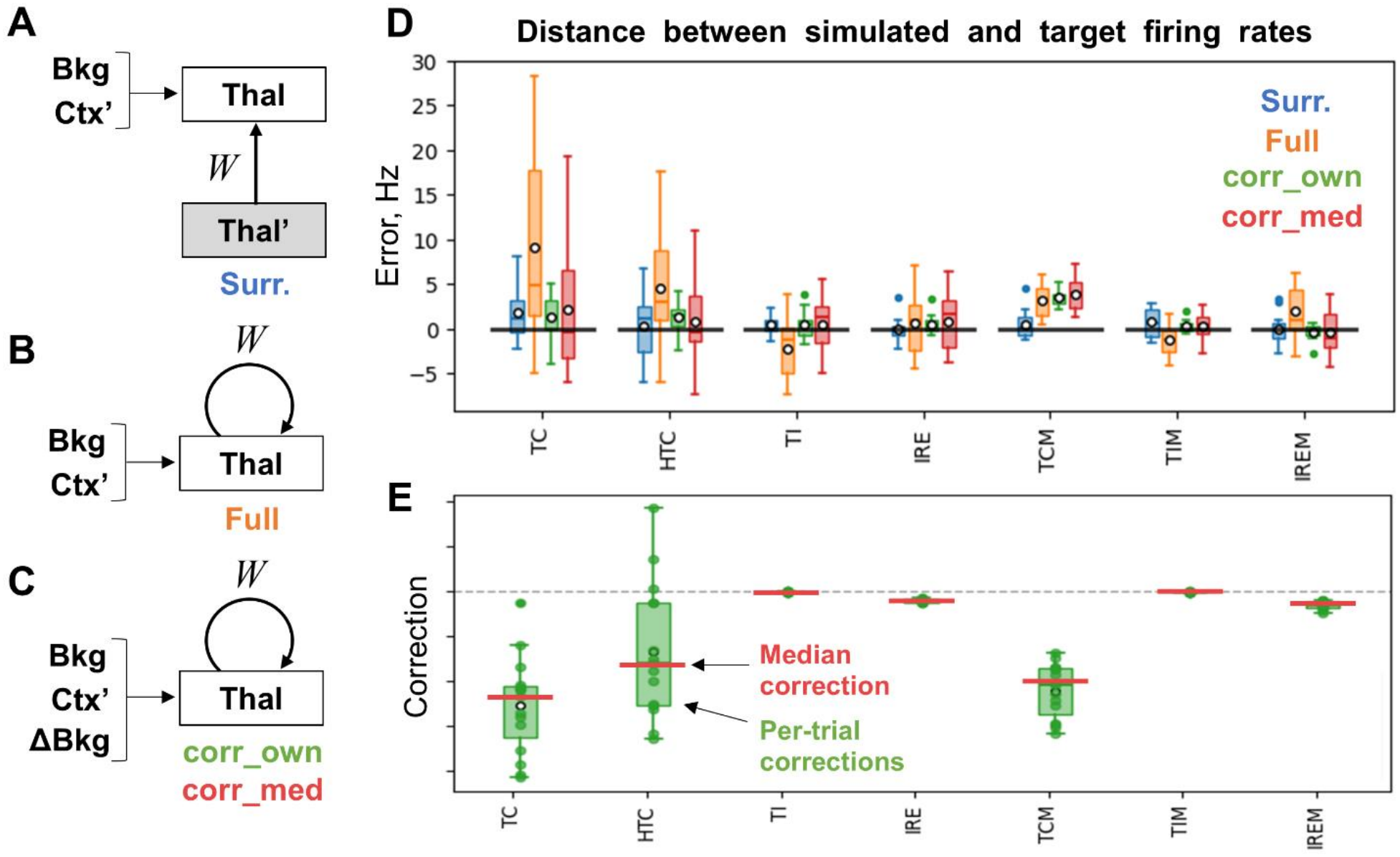
Firing rate errors and secondary tuning in the thalamic subcircuit. **(A)** Schematic of the fully decoupled thalamic subcircuit. Ctx’ and Thal’ – surrogate copies of cortical/thalamic populations, respectively; Bkg – background input. All non-background inputs to Thal originate from surrogate populations. **(B)** Schematic of the recurrently connected thalamic subcircuit with surrogate cortical inputs. Same configuration as in panel B, with additional tonic current corrections (ΔBkg). Two sets of corrections are considered: those optimized separately for each random model realization (corr_own) and the median corrections across realizations (corr_med). **(D)** Distributions of firing rate errors across 15 random model realizations. Blue – fully decoupled thalamic subcircuit (panel A); orange – recurrently connected subcircuit (panel B); green – recurrently connected subcircuit with realization-specific corrections (corr_own); red – recurrently connected subcircuit with median corrections (corr_med). **(E)** Tonic current corrections obtained from the secondary tuning procedure. Green – distributions of corrections optimized separately for each model realization; red horizontal marks – median corrections across realizations. Error definition and boxplot conventions are the same as in Fig. 3C.

We next examined whether gradual fading from surrogate inputs to recurrent connections could help the thalamic subcircuit reach a near-target regime, as it did in the cortex. Freezing either the IREM→TI or IREM→IRE pathway reduced the errors (**Fig. S1**, orange and green bars), but the original error distributions re-emerged in both cases after recurrent connectivity was restored (**Fig. S1**, brown and pink vs. red bars). We also tested the fading procedure starting from the fully decoupled circuit and obtained a similar result (**Fig. S1**, purple vs. red bars). The findings thus provided no evidence for a coexisting near-target regime, suggesting that the errors arose primarily from inaccuracies in surrogate compensation.

#### Second stage of background optimization

To reduce the remaining firing rate errors, we performed a second tuning stage for the thalamic subcircuit. We preserved the background inputs obtained in the fully decoupled setup and introduced seven additional tonic current corrections, one for each thalamic population (**Fig. 5C**). We tuned these corrections jointly based on simulations of the recurrently connected thalamic subcircuit.

To determine the corrections, we used a feedback procedure that continuously adjusted them in response to deviations of the population firing rates from their targets (**Figs. S5 and S6**). We allowed the corrections to settle, then disabled the feedback and continued the simulation with the corrections held fixed. Across 15 random model realizations, the resulting firing rate errors had substantially smaller means and lower variability than in the uncorrected recurrent subcircuit (**Fig. 5D**, green vs. orange bars; **Table S6**, Original vs. Corr. trial).

The optimized corrections varied across realizations (**Fig. 5E**, green bars). To obtain a common set of corrections, we calculated the median correction for each population across the 15 realizations (**Fig. 5E**, red horizontal marks). We then applied these median corrections to each realization and evaluated the resulting firing rate errors. The error distributions were broader than those obtained with realization-specific corrections, but the mean errors remained within ±5 Hz for all thalamic populations (**Fig. 5D**, red vs. green bars; **Table S6**, Corr. trial vs. Corr., median). Thus, the second tuning stage substantially improved the result, demonstrating that the surrogate-based background solution can provide a useful basis for further optimization.

### Thalamocortical circuit

Finally, we tested the tuned parameters in the full thalamocortical model. We used the background inputs obtained in the surrogate open-loop model and re-applied the second-stage corrections previously found for the isolated thalamic subcircuit (**Fig. 5E**). As in the isolated cortical subcircuit, the near-target regime was not reached from naïve initial conditions (**Fig. S2**, orange bars). We therefore used the same surrogate-to-recurrent fading procedure. Initially, all excitatory-to-excitatory connections were replaced by surrogate inputs (**Fig. 6B**, left). After the initial transients decayed, the surrogate inputs were gradually faded out while the corresponding recurrent connections were restored over the course of 2 seconds. The circuit remained near the target regime after recurrent connectivity was fully restored (**Fig. 6B**, right). Population spiking remained largely asynchronous, with irregular low-amplitude fluctuations. An example of firing rate dynamics (for IT3 population) before and after the transition is shown in **Fig. 6C**, and the full traces for all populations are presented in **Figs. S3 and S4**.

**Figure 6.**
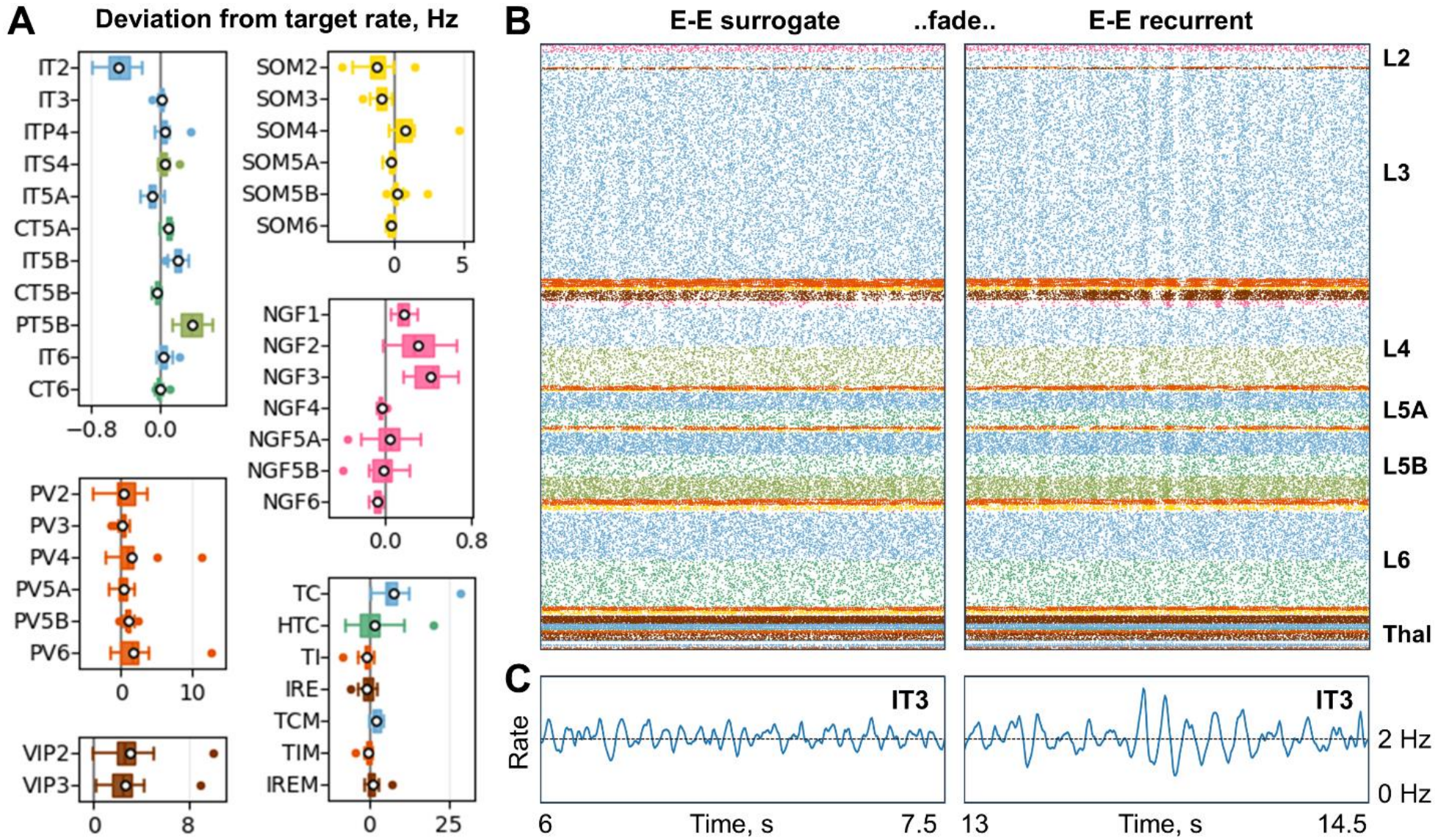
Evaluation of parameter tuning in the full thalamocortical model. Simulations were performed with the background rates obtained in the fully decoupled circuit and the median thalamic current corrections. Recurrent excitatory-to-excitatory (E–E) projections were initially replaced by surrogate inputs and gradually restored between 8 and 10 s. **(A)** Distributions of firing rate errors across 15 random model realizations, calculated from the mean population firing rates over the final 5 s of simulation. Vertical black lines mark zero error. Boxplot conventions are the same as in **Fig. 3C. (B)** Spike rasters from one model realization, shown in two 1.5-s intervals before (left) and after (right) the restoration of recurrent E-E projections. Population colors match those in panel A. **(C)** Firing rate dynamics of the IT3 population in the same time intervals as in panel B. Dashed horizontal lines indicate the target firing rate.

We evaluated the resulting firing rates across 15 random model realizations using the same tuned parameters. Mean firing rate errors remained small for most populations, although the magnitude and variability of the errors differed across them (**Fig. 6A**). Some of the errors were somewhat larger than in the corresponding isolated cortical or thalamic subcircuits, consistent with an effect of thalamocortical and corticothalamic interactions that were absent during the preceding subcircuit tuning stages (**Table S5**, E-E faded; **Table S6**, Corr. median and Isolated vs. Thalamocortical groups). The resulting errors stayed within ±2 Hz for most of the populations, with four populations having errors beyond this range.

## Discussion

In this work, we developed a circuit decoupling method with surrogate compensation for baseline firing rate tuning in recurrent circuit models. We applied it to our 43-population biophysically detailed model of the macaque auditory thalamocortical circuit (Dura-Bernal et al., 2023a). Using full decoupling, we tuned the background inputs for each population separately and reached the target firing rates in the resulting open-loop setup. We then tested the open-loop solution separately in the isolated cortical and thalamic subcircuits with compensated partial decoupling. Both subcircuits deviated substantially from their targets, and our tests suggested different causes. The cortical subcircuit was able to sustain a near-target regime, but with naïve initialization it settled in a coexisting regime far from the target. Gradually fading from surrogate to recurrent self-excitation solved the issue, enabling the subcircuit to reach the near-target regime. Errors in the thalamic subcircuit instead appeared to reflect imperfect surrogate compensation. An additional multidimensional tuning of the subcircuit yielded corrections to the open-loop solution and reduced the errors. Finally, we combined the open-loop background solution, thalamic corrections, and the fading procedure in the full thalamocortical model. Across 15 random model realizations of connectivity and background activity, population firing rates remained close to their targets, with modest deviations in a few populations. Overall, compensated full decoupling enabled background activity to be tuned across populations, while partial decoupling helped diagnose and address the problems that appeared when recurrent connectivity was restored. Together, these methods overcame the challenge of optimizing a large-scale circuit model comprising 43 highly interacting populations of multicompartment biophysical neurons. The approach yielded an improved macaque auditory thalamocortical baseline model with physiological firing rates. This advance enables us, and the broader scientific community, to use the model to investigate auditory cortical function in health and disease.

### Circuit decoupling for background-input tuning

The main benefit of circuit decoupling is the reduction in the number of background-input parameters that must be tuned jointly. Once recurrent interactions are removed, the background inputs to each population can be optimized in a separate low-dimensional search. However, decoupling also omits the recurrent inputs that cells normally receive, so the background inputs tuned in the decoupled setup may yield different firing rates when recurrent connectivity is restored. Our framework and the method proposed by Isbister et al. (2026) address this mismatch at different stages of the tuning process.

Isbister et al. (2026) left the omitted recurrent inputs uncompensated in the decoupled open-loop setup and addressed the potential mismatch in a subsequent stage. They first measured how each population’s open-loop firing rate depended on its background input and then estimated a mapping between population firing rates in the open-loop setup and in the recurrent circuit. To avoid high-dimensional sampling, they restricted the target firing rate vectors for the recurrent circuit to scalar multiples of a fixed reference vector. Because the corresponding open-loop rates were unknown, they refined the mapping iteratively, using the firing rates obtained in each round of recurrent-model simulations to select candidate background inputs for the next round. In their model, the target rates were reached after a few iterations, although the study did not establish general conditions for convergence.

Our approach instead uses surrogate spike trains to compensate for omitted recurrent inputs in the open-loop setup. These spike trains are generated at the target firing rates of the corresponding presynaptic populations, independently of ongoing circuit activity, allowing background inputs to be tuned separately for each population. Thus, the main distinction between the two methods is how they account for recurrent interactions: our method approximates their effects during open-loop tuning, whereas Isbister et al. estimate the mapping between open-loop and recurrent firing rates through subsequent simulations. In principle, the two approaches could also be combined: compensated open-loop tuning applied to several multiples of the target rate vector could provide initial estimates for the iterative procedure of Isbister et al., potentially improving convergence.

Our framework also supports partial decoupling, in which selected recurrent projections are replaced with surrogate inputs while the remaining connections are preserved. One application is to isolate a subcircuit from the rest of the network while preserving its internal recurrent connectivity. This enables separate analysis of individual subcircuits and limits the scope of joint tuning when it is required. Billeh et al. (2020) and Dura-Bernal et al. (2023a) used a related strategy to tune synaptic weights, progressing from individual layers to the full network. Another application of partial decoupling is to freeze specific recurrent projections to examine their influence on the circuit dynamics. Gradually restoring frozen projections can also help the circuit reach its target when simulations initiated with full connectivity settle into a co-existing off-target regime. Together, these capabilities extend circuit decoupling beyond independent background-input tuning to the analysis and refinement of recurrent circuit activity.

### Conditions for accurate surrogate compensation

Background inputs tuned in the open-loop setup are intended to yield a near-target regime when recurrent connectivity is restored. The success of this transfer depends on how closely the surrogate inputs match the recurrent synaptic inputs that emerge near the target regime. In our implementation, all units within a surrogate population generated independent Poisson spike trains at the same rate. This construction simplifies two aspects of recurrent activity: spike-train correlations and within-population heterogeneity of firing rates. The accuracy of surrogate compensation therefore depends on how strongly these simplifications alter the inputs received by postsynaptic cells.

Replacing each presynaptic cell with an independent Poisson generator removes both temporal correlations within individual spike trains and correlations between cells. This simplification is broadly consistent with experimental observations of irregular spiking in awake cortex and weak pairwise correlations in both cortex and thalamus (Compte et al., 2003; Maimon and Assad, 2009; Ecker et al., 2010; Erisken et al., 2014; Maksimov et al., 2018). Such activity is also well studied theoretically and commonly emerges in large recurrent networks through excitation-inhibition balance and other related mechanisms (van Vreeswijk and Sompolinsky, 1998; Brunel, 2000; Renart et al., 2010; Sanzeni et al., 2022). Together, these findings support the plausibility of the independent Poisson approximation for the chosen baseline regime.

The other simplification concerns cell-to-cell firing rate heterogeneity, which is absent from our surrogate populations whose units all fire at the same rate. By comparison, experiments show that spontaneous firing rates vary widely across cells in the cortex (Hromádka et al., 2008; O’Connor et al., 2010) and thalamus (Anderson and Linden, 2011; Bartlett and Wang, 2011). Broad rate distributions also emerge naturally in network models with random or heterogeneous connectivity (van Vreeswijk and Sompolinsky, 1998; Roxin et al., 2011; Vegué and Roxin, 2019; Sanzeni et al., 2022). Thus, equal-rate surrogates do not reproduce this well-established aspect of neuronal activity, which may reduce the accuracy of surrogate-based tuning.

The negative impact of presynaptic rate averaging depends on the resulting distortion of the effective input to postsynaptic cells. This distortion is expected to remain small under two main conditions. First, the mean postsynaptic effect must depend approximately linearly on presynaptic firing rate, while strong synaptic non-linearities could violate this condition (Mongillo et al., 2012). Second, each postsynaptic cell must receive input from a sufficiently large and unbiased sample of presynaptic cells, so that the mean firing rate in that sample stays close to the presynaptic population mean (Vegué and Roxin, 2019). Violations of either condition could reduce the accuracy of surrogate compensation and decrease tuning quality.

### Achieving a near-target regime in recurrent circuits

Firing rate errors emerging after restoring recurrent connectivity do not necessarily reflect inaccurate background-input tuning. Several regimes may co-exist in a recurrent circuit, and the regime it reaches can depend on the initial conditions (Schmidt et al., 2018). Thus, even when a tuned circuit can support a near-target regime, a naïvely initialized simulation may settle into a different co-existing regime with substantially larger errors. Distinguishing between these possibilities is important for determining whether additional background tuning is needed. We used surrogate-compensated partial decoupling to examine this question in the isolated cortical and thalamic subcircuits.

The cortical subcircuit illustrated the importance of proper initialization. With all intracortical connections recurrent from the simulation outset, the circuit settled into a regime with large firing rate errors. However, when we started with surrogate self-excitation and then gradually restored the original recurrent connectivity, the circuit successfully maintained a near-target regime (**Fig. 4C**). Thus, our framework provided a way to reach the required regime even when multiple stable regimes co-exist.

The thalamic subcircuit presented a different problem. Gradual restoration of recurrent connections, starting from either partially or fully decoupled configurations, led to the same errors as under naïve initialization (**Fig. S1**). These results provided no evidence for a coexisting near-target regime and instead suggested that the errors arose primarily from inaccurate surrogate compensation. The thalamic results therefore pointed to a need for additional tuning.

Addressing the thalamic errors required a procedure that accounts for interactions between the populations. We introduced an additional input correction for each thalamic population and tuned these corrections jointly, based on simulations of the recurrently connected thalamic subcircuit. This strategy helped us substantially reduce the errors (**Fig. 5D**) but reintroduced multidimensional search, diminishing the computational advantage of independent population-wise tuning. However, our use of compensated partial decoupling kept the joint search confined to the seven thalamic populations, avoiding 43-population full-circuit optimization. The background inputs obtained in the fully decoupled setup may also have simplified the search by providing a starting point closer to the target regime than randomly chosen inputs would yield. Thus, even when independent population-wise tuning was insufficient, our approach could facilitate further optimization by restricting its scope and providing a suitable starting point.

The complete thalamocortical circuit provided a final test of the staged tuning approach, including the effects of interactions between the separately tuned subcircuits. With the background inputs and thalamic corrections obtained in the preceding stages, and using the previously established fading procedure, we achieved firing rates close to their targets in most populations (**Fig. 6A**). Some errors increased relative to the isolated subcircuits, consistent with the influence of interactions between cortex and thalamus that had previously been approximated by surrogate inputs. Overall, these results demonstrate that compensated decoupling can support staged baseline tuning in a large recurrent circuit without requiring joint optimization across all populations.

### Toolbox and future extensions

We implemented our method as a toolbox that automatically modifies NetPyNE-compatible model descriptions. The toolbox applies circuit decoupling with surrogate compensation based on user-defined rules that specify which connections should be replaced by surrogate inputs and which populations should be omitted. The toolbox supports full decoupling for independent background-input tuning, compensated isolation of selected recurrent subcircuits, and selective connection replacement for pathway freezing or gradual surrogate-to-recurrent fading.

Although we demonstrated these capabilities in one model and for one target regime, the toolbox is potentially applicable to a broad class of NetPyNE-compatible circuit models. More than 100 such models are publicly available (EBRAINS, n.d.), spanning diverse brain regions, functions, and diseases and contributing to over 40 publications (e.g., Medlock et al., 2022; Dura-Bernal et al., 2023a,b; Ponzi et al., 2023). The method could also be adapted to other modeling frameworks, such as BMTK (Dai et al., 2020).

Future extensions could improve the match between surrogate inputs and recurrent activity. One approach would be to iteratively refine the surrogate inputs to better reproduce the firing rate distributions and spike-train statistics observed in the recurrent network. After each recurrent-network simulation, these activity properties could be measured and used to construct updated surrogates for the next round of open-loop tuning, with the process repeated until convergence. Although this procedure would still ignore cross-cell correlations, it could further reduce errors caused by surrogate-recurrent mismatch.

Another promising future direction is to extend the present surrogate-compensated decoupling framework from stationary firing rate targets to time-dependent activity, such as evoked responses or oscillations. For such applications, surrogate inputs would need to capture the relevant temporal structure in addition to mean firing rates. Although we did not test this extension here, it could broaden the use of circuit decoupling from establishing baseline regimes to tuning and analyzing dynamic properties of biophysically detailed circuit models.

## Methods

### Model

We used a modified version of the biophysically detailed model of the macaque auditory thalamocortical circuit developed by Dura-Bernal et al. (2023a). Most model parameters were retained from the published version. Below, we briefly describe the model and the modifications introduced for the present study.

### Cells and populations

The model contained 12,908 neurons in 43 populations, including 36 cortical populations across layers 1-6 (including L5A/L5B subdivision) and 7 populations in the auditory thalamus (medial geniculate body, MGB) and thalamic reticular nucleus. Cortical excitatory cells belonged to the types: L2-6 intratelencephalic pyramidal (IT), L4 stellate (ITS4), L5-6 corticothalamic (CT), and pyramidal-tract-like cells of L5B (PT5B). Inhibitory cell types were: parvalbumin-positive (PV), somatostatin-positive (SOM), vasoactive intestinal peptide-positive (VIP), and neurogliaform (NGF). L1 contained only NGF cells; L2-6 contained all inhibitory types. The thalamic part contained MGB thalamocortical cells (TC, HTC, TCM), MGB interneurons (TI, TIM), and reticular nucleus cells (IRE, IREM); it was divided into the core (TC, HTC, TI, IRE) and matrix (TCM, TIM, IREM). Cells were randomly positioned within their respective layers based on experimentally estimated densities.

Cell models contained 1-6 compartments (depending on the type) whose membrane properties were defined by several types of ion channels with biologically realistic parameters. Some of the cellular parameters were changed compared to the published model, summarized in **Table S8**. These changes were made for (1) thalamocortical cells to achieve more realistic single-cell activity and (2) excitatory cortical cells to avoid cell-level multistability.

### Connectivity

The circuit had random sparse connectivity with the same connection probabilities and synaptic delays as in Dura-Bernal et al. (2023a). Most of the synaptic properties remained the same, except for revised time constants and reversal potential of GABAB receptors; also, some of the pathways had a different GABAA:GABAB ratio in our version (see **Table S8**). Furthermore, we revised the synaptic positioning rules that define pathway-specific lists of postsynaptic cell sections to choose from. As in the published model, every section contained one numerical segment, so section choice fully specified synapse location at the resolution of the used cell models. The complete section map is given in **Table S9**.

The main connectivity changes involved synaptic weights. Dura-Bernal et al. (2023a) combined the available experimental evidence to build the original weight matrix and introduced a set of multipliers jointly applied to this matrix that were tuned to fit the required circuit activity. In this paper, we tuned the background inputs rather than weights, so we removed the multipliers used by Dura-Bernal et al. (2023a). Instead, we applied two non-specific multipliers: 0.125 to the connections between excitatory cells (E-E) and 0.25 to the other connections. The weights of E-E connections were reduced compared to the others to prevent abnormally synchronized circuit activity.

### Background inputs

Every neuron in our model received a background input consisting of two independently generated Poisson spike trains delivered to the soma. One spike train provided excitatory input through an AMPA synapse, and the other – inhibitory input through a GABAA synapse. Neurons in the same population shared the two input rates and synaptic weights, but received independently generated spike trains. The synaptic weights were chosen separately for each population so that an individual background spike (from either excitatory or inhibitory input) received in the resting state (with all other synaptic inputs turned off) produced a somatic voltage deflection with the absolute amplitude of approximately 0.5 mV (see *w*_*xe*_ and *w*_*xi*_ columns in **Table S3**). We chose this value arbitrarily from a biologically plausible EPSP/IPSP range and used it for each cell for simplicity. Such a choice is reasonable due to the abstract nature of our background inputs that imitate the joint influence of processes not directly accounted for in the model, including long-range inputs from other regions and neuromodulatory inputs. The background weights were kept constant, while the background rates (*r*_*xe*_, *r*_*xi*_) were the tunable variables. All the background inputs used in the published version of the model to provide minimal baseline activity were removed.

For the thalamic and excitatory cortical populations, we also added a tonic offset current to the soma intended to achieve a biologically plausible resting state (**Table S2**).

### Simulation and analysis

We configured and ran simulations using NetPyNE (Dura-Bernal et al., 2019), with NEURON as the underlying simulation engine. Simulations ran on a computing cluster, with each network distributed across 1-64 CPU cores. We used a fixed integration time step of 0.05 ms and a model temperature of 37 °C. Recurrent connectivity, surrogate inputs, and background spike trains were governed by separate pseudo-random generators. For validation, secondary thalamic tuning, and comparisons of model configurations, we ran 15 trials per condition, with different random seeds. Within each condition, background-input rates and synaptic parameters remained fixed, while the connectivity and input spike trains were resampled in every trial.

We calculated each population’s mean firing rate by dividing the spike count within the analysis interval by the interval duration in seconds and the population size, including silent neurons. The coarse background-input scan used 5-s simulations and a 35 s analysis interval; the fine scan used 12-s simulations and a 7–12 s interval. Validation of the selected background inputs and comparisons of isolated cortical and thalamic subcircuits used 10-s simulations and a 7–10 s interval. Secondary thalamic tuning and its long-duration controls used 220-s simulations and a 200–220 s interval. The final full-circuit evaluation used 20-s simulations and a 15–20 s interval.

We calculated firing rate errors by subtracting each population’s target rate from its mean firing rate in each trial. **Tables S5 and S6** report the mean error ± sample standard deviation across realizations, in Hz. Boxplots show medians, interquartile ranges, and means (white circles); whiskers extend to the most extreme values within 1.5 interquartile ranges of the box edges, with outliers plotted individually. Illustrative rasters and firing rate traces represent individual trials. For **Figs. 2D, 6C, S3, and S4**, we calculated instantaneous population firing rates in 5-ms bins and smoothed the traces using a Gaussian kernel with a standard deviation of 5 ms.

### Circuit decoupling with surrogate compensation

We decoupled individual populations or subcircuits by replacing selected synaptic inputs from neurons of the original model with inputs from Poisson spike generators (**Fig. 1**). For each original population *P*_*m*_, we constructed a surrogate population 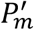 consisting of Poisson generators distributed within the same spatial region and at the same density as the neurons of *P*_*m*_. Each generator emitted an independent, homogeneous Poisson spike train at the target mean firing rate 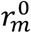 that we specified for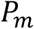.

### Full decoupling for isolating populations

To fully decouple the circuit, we replaced every projection *P*_*m*_ → *P*_*n*_ with a corresponding surrogate input 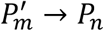. This included self-projections, which comprise synaptic connections between neurons of the same population: each self-projection *P*_*m*_ → *P*_*m*_ was replaced with a surrogate input 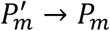. We generated the surrogate connectivity using the same rules as for the original recurrent connectivity, preserving connection probabilities, synaptic-weight distributions, transmission delays, and distance-dependent terms; we also preserved the synaptic mechanisms and postsynaptic placement rules. **Figure 1A,B** illustrates this construction for two example populations *P*_1_ and *P*_2_ and their corresponding surrogate populations 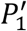 and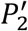 . With all recurrent connections replaced by fixed-rate surrogate inputs, changing one population’s background input affected only that population’s activity, allowing the background inputs to be tuned independently for each population.

### Partial decoupling for subcircuit isolation and connection freezing

For partial decoupling, we selected a subset of recurrent projections and replaced them with surrogate inputs while keeping the other recurrent projections intact. We refer to this replacement of a selected projection as freezing. The middle panel of **Fig. 1C** illustrates this operation for the *P*_1_ → *P*_3_ projection, which is replaced by the surrogate input 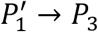, while the other projections among *P*_1_, *P*_2_, *P*_3_ remain unchanged.

To isolate a subcircuit comprising a set of populations *S*, we retained all recurrent projections between populations in *S*, including their self-projections, and replaced every projection into *S* from an outside population with a surrogate input. The right panel of **Fig. 1C** illustrates this procedure for a subcircuit *S* = {*P*_2_, *P*_3_}: the recurrent projections between *P*_2_ and *P*_3_ and their self-projections are preserved, while the projections *P*_1_ → *P*_2_ and *P*_1_ → *P*_3_ originating outside *S* are replaced by the surrogate inputs 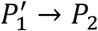 and 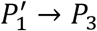, respectively.

### Semi-surrogate connections

Our method also allowed creating semi-surrogate projections, in which synaptic input is divided between recurrent and surrogate sources (**Fig. 1D**). For each selected projection *P*_*m*_ → *P*_*n*_, we retained the original recurrent connections and independently generated a surrogate projection 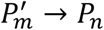 using the same connection rules and a different random seed. The two projections supplied input to the same postsynaptic population but had separately sampled connections and different sources.

We controlled the relative contributions of the two sources using a splitting coefficient *k ∈* [0,1]. The effective synaptic weights of the recurrent and surrogate inputs were given by:

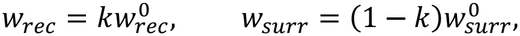

where 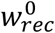 and 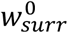 denote the unscaled weights of synapses in the recurrent and surrogate projections, respectively. Thus, *k* = 0 corresponds to fully surrogate input, and *k* = 1 corresponds to the original recurrent projection.

### Background input tuning in a fully decoupled circuit

We tuned the background inputs to our 43-population thalamocortical circuit to achieve the target regime *R*_0_ specified as a vector of *N* = 43 population target firing rates: 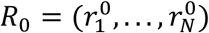,which we chose based on the literature (**Table S1**). We started the background-input tuning process by using a fully decoupled circuit with surrogate compensation (**Fig. 2A**). All the recurrent connections of the original circuit A1 were replaced by compensatory inputs from a surrogate circuit A1^′^ comprised of fixed-rate Poisson spike generators.

For each population *P*, we performed a two-stage search of the background-rate pair 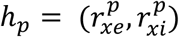 : a coarse rectangular grid scan followed by a fine grid scan within a parallelogram fitted around the region where the coarse scan yielded firing rates near the target (**Fig. 2**). Both scans used an *M* × *M* grid (*M* = 20), indexed by *m*_*e*_, *m*_*i*_ = 1, …, *M*.

### Coarse scan

For the coarse scan, we sampled the background rates for a population *P* uniformly from pre-defined intervals:

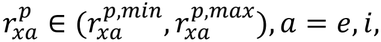

where 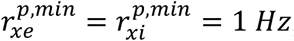, and 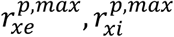 were chosen for each population separately (**Table S7**).

We simulated multiple populations at once for each grid point, with the population-specific background inputs defined by the grid coordinates (*m*_*e*_, *m*_*i*_):

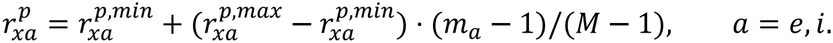

The populations shared the same surrogate inputs and did not affect each other due to decoupling.

We ran each simulation for 5 s and measured mean population firing rates over 3–5 s interval. The measurements at all grid points formed the coarse firing rate map 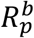 for each population *P* (**Fig. 2B**).

### Fitting the fine-search region

For each population *P*, we fitted a parallelogram *Q*_*p*_ encompassing a region on the 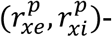 plane where the coarse firing rate map 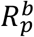 had values close to the target rate 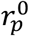.

The procedure of *Q*_*p*_ fitting was based on finding low-rate and high-rate contours on the coarse map 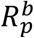. For the low-rate level, we used 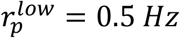 for every population. For the high-rate level 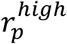 we used a predefined rate of 10, 30, or 60 Hz for cortical populations and the target rate 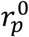 for thalamic populations (**Table S7**).

For the cortical populations, we extracted the contours directly from the coarse map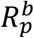 . For each contour, we chose the segment containing the point with the smallest 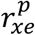 value. For the thalamic populations, we first applied Gaussian smoothing to the map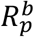, with a standard deviation of one coarse-grid cell per axis, and retained the longest segment for each contour.

We approximated the high-rate contour by a straight line through the leftmost and rightmost points and computed its slope *s*_*p*_. We then calculated the maximum horizontal gap *d*_*p*_ between this line and the low-rate contour. We defined a diagonal strip bounded by two lines parallel to the fitted line, using a width-expansion fraction *α*_*p*_ and a right-edge shift fraction *β*_*p*_ (**Table S7**). We set the horizontal width to (1 + *α*_*p*_)*d*_*p*_ and obtained the right edge by shifting the fitted line rightward by *β*_*p*_*d*_*p*_.

We then limited the strip along the 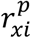 -axis to form the search parallelogram *Q*_*p*_. We set its lower edge at 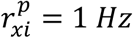 and its upper edge at the same inhibitory rate as the rightmost vertex of the original high-rate contour. If the lower-right corner had 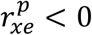, we raised the lower edge until 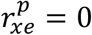 at that corner. We rounded the lower-right corner coordinates, horizontal width, and vertical height to multiples of 10 Hz. We used these four values and the fitted slope *s*_*p*_ to define the fine grid (**Table S7**).

### Fine scan

For the fine scan, we used the indices *m*_*e*_, *m*_*i*_ = 1, …, *M* to specify positions on a skewed grid covering each population’s fitted parallelogram *Q*_*p*_. We denote its lower-right corner by 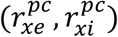, and its horizontal width and vertical height by 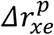 and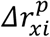, respectively (**Table S7**). Using the fitted slope *s*_*p*_, we calculated the background rates at the grid nodes:

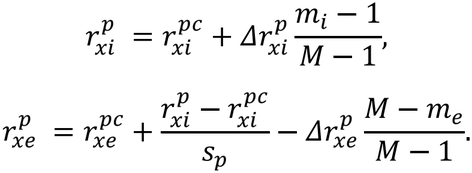

The index *m*_*i*_ specified rows at uniformly spaced inhibitory rates 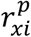 from the lower to the upper bound of *Q*_*p*_. Within each row, *m*_*e*_ specified uniformly spaced excitatory rates 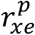 from the left to the right diagonal edge of *Q*_*p*_. For each grid point, we simulated multiple populations together, assigning each population the background rates from its own parallelogram.

Some parallelograms extended into the region of negative excitatory background rates. For each affected population, we skipped simulations for grid points with negative excitatory rates and marked the corresponding entries of the population’s firing rate map as missing. We ran each simulation for 12 s and measured mean population firing rates over 7-12 s. These measurements formed the fine firing rate map *R*_*p*_ for each population (**Fig. 2C**).

### Selecting the background rates

We linearly interpolated each fine firing rate map *R*_*p*_ onto a 200 × 200 grid in the grid-index coordinates (*m*_*e*_, *m*_*i*_). In these coordinates, *Q*_*p*_ is represented as a rectangle (**Fig. 2C**). We applied Gaussian smoothing with a standard deviation of 6 interpolation-grid cells per axis for cortical populations and approximately 11.7 cells per axis for thalamic populations.

From the smoothed map, we extracted the contour at the target firing rate 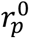 and selected its middle point by index (**Fig. 2C**). We converted the (*m*_*e*_, *m*_*i*_) grid coordinates of the selected midpoint to background-rate coordinates 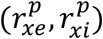 and rounded them to the nearest integer value in Hz. We denote the resulting background-rate pair for population *P* by 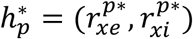 (**Figure 2C,D**) and the vector of such pairs for all populations by 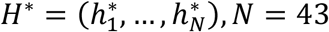. The results of this process are given in **Table S3**.

### Validation across random realizations

With *H*^*\**^ fixed, we ran 15 simulations of the fully decoupled circuit for 10 s each, resampling connectivity and Poisson spike trains independently for each trial, and calculated the mean firing rates over 7-10 s interval. For each population and trial, we calculated the firing rate error by subtracting the target rate 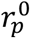 from the measured mean rate. The resulting error distributions for all cortical and thalamic populations are represented by blue boxplots in **Figs. 3C and 5D**, respectively, and the summary statistics are given in **Tables S5 and S6**.

### Fading from surrogate to recurrent inputs

We used semi-surrogate projections to implement gradual transitions from surrogate to recurrent connectivity, a procedure we refer to as fading. For each selected projection, we varied the splitting coefficient *k*(*t*) from 0 to 1, increasing it linearly over a transition interval (*t*_*a*_, *t*_*b*_):

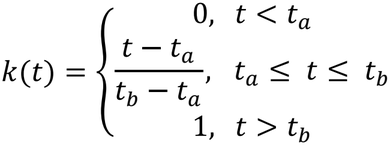

In our NEURON implementation, we scaled the instantaneous synaptic conductances of the recurrent and surrogate projections by *k*(*t*) and 1 − *k*(*t*), respectively, while keeping the connection weights, synaptic properties, and surrogate firing rates fixed. All affected connections within each simulation rank shared references to the corresponding multipliers. Thus, fading required updating only these shared values at each simulation time step, rather than modifying individual connections. We applied the same coefficient to all selected projections and all receptor components within each projection.

In the isolated cortical subcircuit, we faded all excitatory-to-excitatory (E-E) projections from surrogate to recurrent input over 3–5 s (**Fig. 4**). In the isolated thalamic subcircuit, we tested fading of all intrathalamic projections and of the individual inhibitory projections IREM→TI and IREM→IRE, in separate experiments, using the same transition interval as in the cortical subcircuit (**Fig. S1**). In the complete thalamocortical circuit, we faded all existing projections between excitatory populations, including cortical and thalamic populations, over 8–10 s (**Fig. 6**).

### Secondary tuning stage

For the second tuning stage, we introduced an additional background-input correction for each thalamic population, implemented as a current injected to the soma. We used an optimization procedure based on feedback control of the firing rates to jointly tune the thalamic corrections. With active feedback, the controller dynamically adjusted the correction currents to bring population firing rates toward their targets. To obtain corrections that could maintain near-target firing rates without ongoing feedback, we split the feedback signal into slow and fast components and accumulated the corrections produced by the slow component. After allowing the controlled activity to settle, we disabled the feedback and fixed the slow corrections, discarding the transient components.

### Population firing rate estimate

For population *P* containing *N*_*p*_ neurons, we defined the normalized population spike train as:

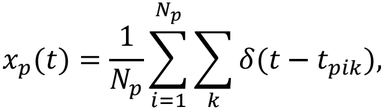

where *t*_*pik*_ is the time at which spike *k* from neuron *i* reached the controller, including a delivery delay of 1 *ms*. We estimated the running mean firing rate *r*_*p*_(*t*) by exponential filtering with time constant *τ*_*r*_:

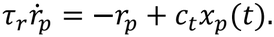

We used the conversion factor *c*_*t*_ = 10^3^ *ms s*^−1^ to express the rate estimate in hertz while *t* is given in milliseconds. Each delivered spike increased *r*_*p*_ by *c*_*t*_/(*N*_*p*_*τ*_*r*_), and between spikes *r*_*p*_ decayed exponentially with time constant *τ*_*r*_. Normalization by *N*_*p*_ yielded the mean firing rate per neuron.

### Feedback control of tonic current

We defined the firing rate error as *e*_*p*_(*t*) = *r*_*p*0_ − *r*_*p*_(*t*), where *r*_*p*0_ is the target rate of population *P*. We filtered the error signal with time constant *τ*_*m*_ to obtain its slowly varying component *m*_*p*_:

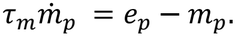

We calculated the slow feedback signal *s*_*p*_ based on *m*_*p*_, the fast feedback signal *u*_*p*_ based on the residual fast error component *e*_*p*_ − *m*_*p*_, and the total feedback signal *z*_*p*_ as their sum:

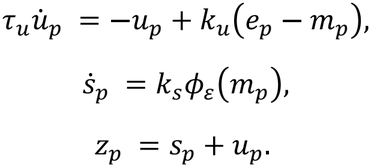

Here, *τ*_*u*_ and *k*_*u*_ are the time constant and the gain of *u*_*p*_, respectively; *k*_*s*_ is the gain of *s*_*p*_; *ϕ*_*ε*_ is a dead-zone function. The full list of parameter values is given in **Table S10**.

We limited the contribution of the fast feedback *u*_*p*_ by choosing a small value for its gain *k*_*u*_. This allowed the slow branch *s*_*p*_ to accumulate the correction for a persistent firing rate mismatch without relying on the fast dynamic feedback created by *u*_*p*_. We additionally limited the sensitivity of *s*_*p*_ to small firing rate fluctuations by introducing a dead zone:

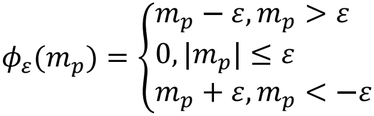

The slow correction *s*_*p*_ therefore accumulated only the portion of the filtered error that exceeded the dead zone.

To close the feedback loop, we used the controller output *z*_*p*_ to drive a somatic current injected to every cell of the controlled population *P*:

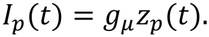

When the population activity *r*_*p*_(*t*) was below the target rate *r*_*p*0_, the error *e*_*p*_(*t*) = *r*_*p*0_ − *r*_*p*_(*t*) was positive, as was its filtered version *m*_*p*_. With *m*_*p*_ > *ε*, the feedback current *I*_*p*_ changed in the depolarizing direction, tending to increase *r*_*p*_(*t*). Conversely, when *r*_*p*_(*t*) was above *r*_*p*0_, the feedback current changed towards hyperpolarization. Thus, the feedback controller worked to keep the activity close to the target *r*_*p*0_. The gain *g*_*µ*_ was chosen to balance the convergence time and stability of the system.

We implemented the controller equations as a NEURON mechanism that received every output spike from the controlled population *P* with the weight 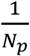. The mechanism’s output drove a current delivered to the soma of every neuron in *P*. In distributed simulations, each MPI rank had its own instance of the controller mechanism. Each instance received spikes from all neurons in *P*, including those on other ranks, and supplied the same control signal to the neurons located on its own rank.

### Tuning protocol and static corrections

To determine the corrections, we simulated the isolated thalamic subcircuit with surrogate cortical input for 220 seconds. We initialized the rate estimate and controller states as *r*_*p*_ = *m*_*p*_ = *u*_*p*_ = 0 and *s*_*p*_ = *z*_0_ = 0 *Hz*, where *z*_0_ is the initial controller output. During the first 10 *s*, only the rate estimate evolved, and the controller output remained at *z*_0_. The feedback was turned on at *t*_0_ = 10 *s* and continued until *t*_*lock*_ = 160 *s*. At locking, we froze the controller states and removed the transient correction, leaving the resulting slow correction as the tonic output:

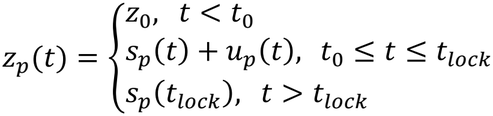

We repeated the tuning procedure across 15 trials with random model realizations. For realization *j*, the fixed current correction for population *P* was:

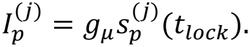

Finally, we obtained one correction per population by taking the median over model realizations. A summary of the results is presented in **Table S4** and **Fig. 5E**. Full dynamics of the controller variables are shown in **Figs. S5 and S6**.

## Acknowledgements

This work was supported by NIH grants R01MH134118, R01DC012947, R01DC019979, P50MH109429, and RF1NS133972; ARL Cooperative Agreement W911NF-22-2-0139; and NYS DOH SCIRB award C38328GG.

## Notes

### Competing Interest Statement

The authors have declared no competing interest.

