## Supplementary figures and tables for "Circuit decoupling with surrogate compensation for baseline tuning in biophysically detailed neural models"

### Figures

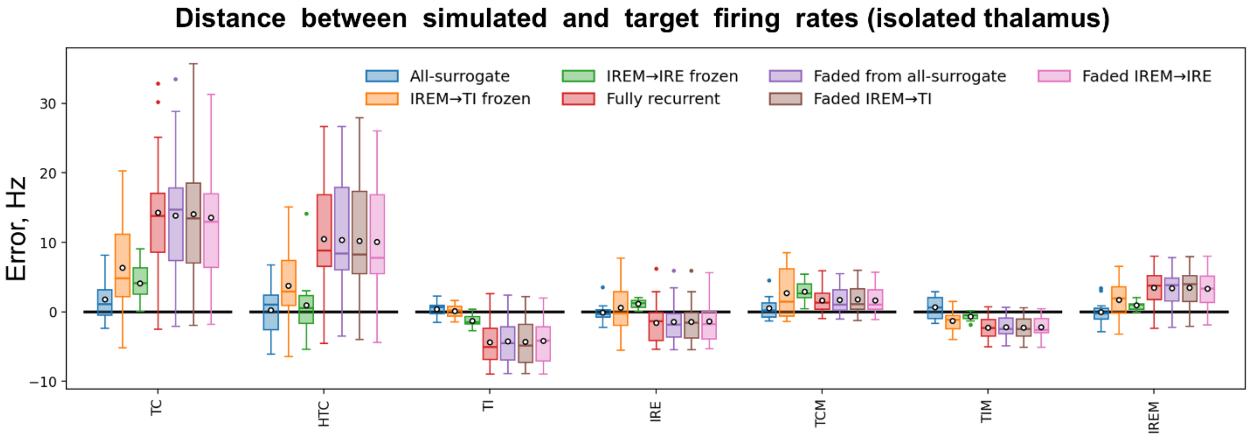


**Figure S1. Surrogate-to-recurrent connection fading in the isolated thalamic circuit.** Distributions of firing rate errors (simulated mean rate minus target rate) are shown for seven conditions: blue – fully decoupled circuit with all-surrogate input; orange – recurrently connected circuit with frozen IREM→TI projection; green – recurrently connected circuit with frozen IREM→IRE projection; red – fully connected circuit; purple – endpoint after fading from the all-surrogate condition; brown – endpoint after fading from the IREM→TI-frozen condition; pink – endpoint after fading from the IREM→IRE-frozen condition. Frozen projections are replaced by fixed-rate surrogate inputs; fading gradually restores their recurrent connectivity over 3–5 s. All simulations are performed without thalamic current correction. Rates were averaged over the 7–10 s interval. Simulations were repeated with 15 random seeds per condition. Boxes – IQR, horizontal lines – medians, white circles – means, whiskers – most extreme samples within 1.5 IQR of the box edges, colored dots – outliers.

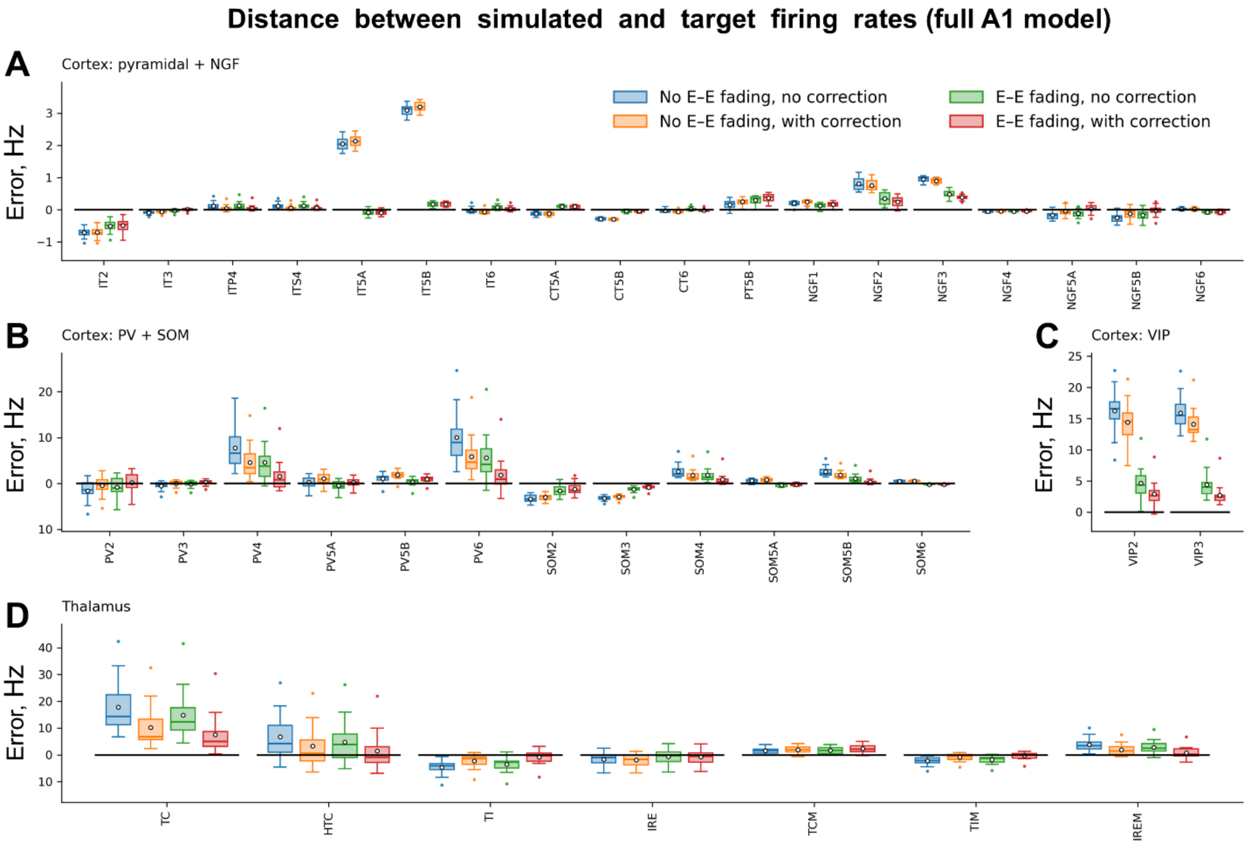


**Figure S2. Effects of excitatory-to-excitatory (E-E) fading and thalamic current correction on population firing rates in the full A1 model.** Population firing rate errors (observed minus target) are shown for cortical pyramidal and NGF populations **(A)**, PV and SOM interneurons **(B)**, VIP interneurons **(C)**, and thalamic populations **(D)**. Colors indicate simulations without either intervention (blue), with thalamic correction alone (orange), with E-E fading alone (green), or with both (red). Rates were averaged over 10-15 s interval. Simulations were repeated with 15 random seeds per condition. Boxes – IQR, horizontal lines – medians, white circles – means, whiskers – samples within 1.5 IQR, colored dots – outliers.

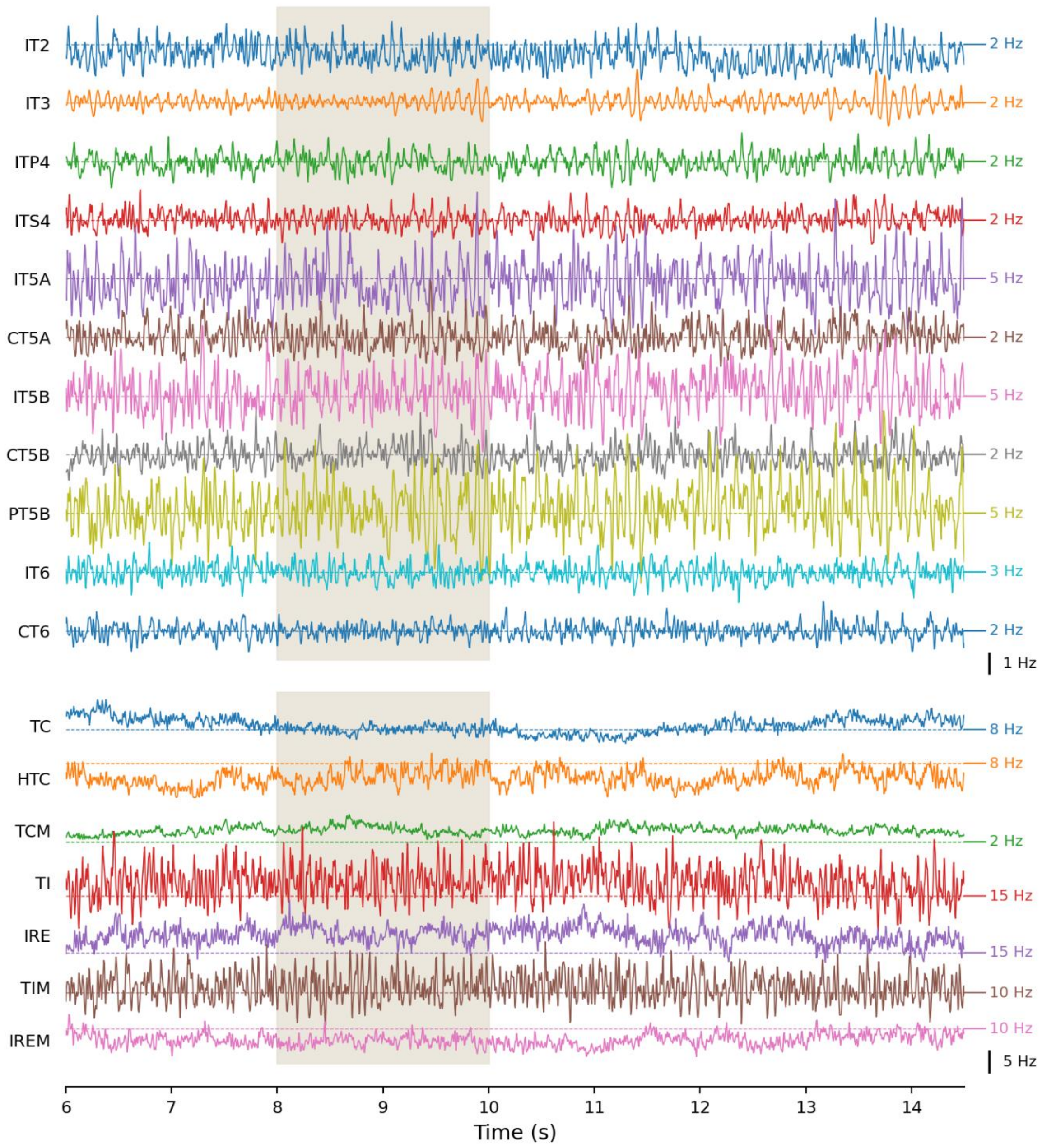


**Figure S3. Firing rate dynamics in the full thalamocortical model (thalamic and excitatory cortical populations).** Each trace shows the mean firing rate in one population, calculated in 5-ms bins and smoothed with a Gaussian kernel (std.: 5 ms). Colored dashed lines indicate target firing rates. Traces are vertically offset, with displacement from the corresponding dashed line representing deviation from target. Black scale bars apply to all traces within each panel. The same model realization is shown as in **Fig. 6B,C**. Grey shading marks the interval of fading from surrogate to recurrent excitatory-to-excitatory inputs.

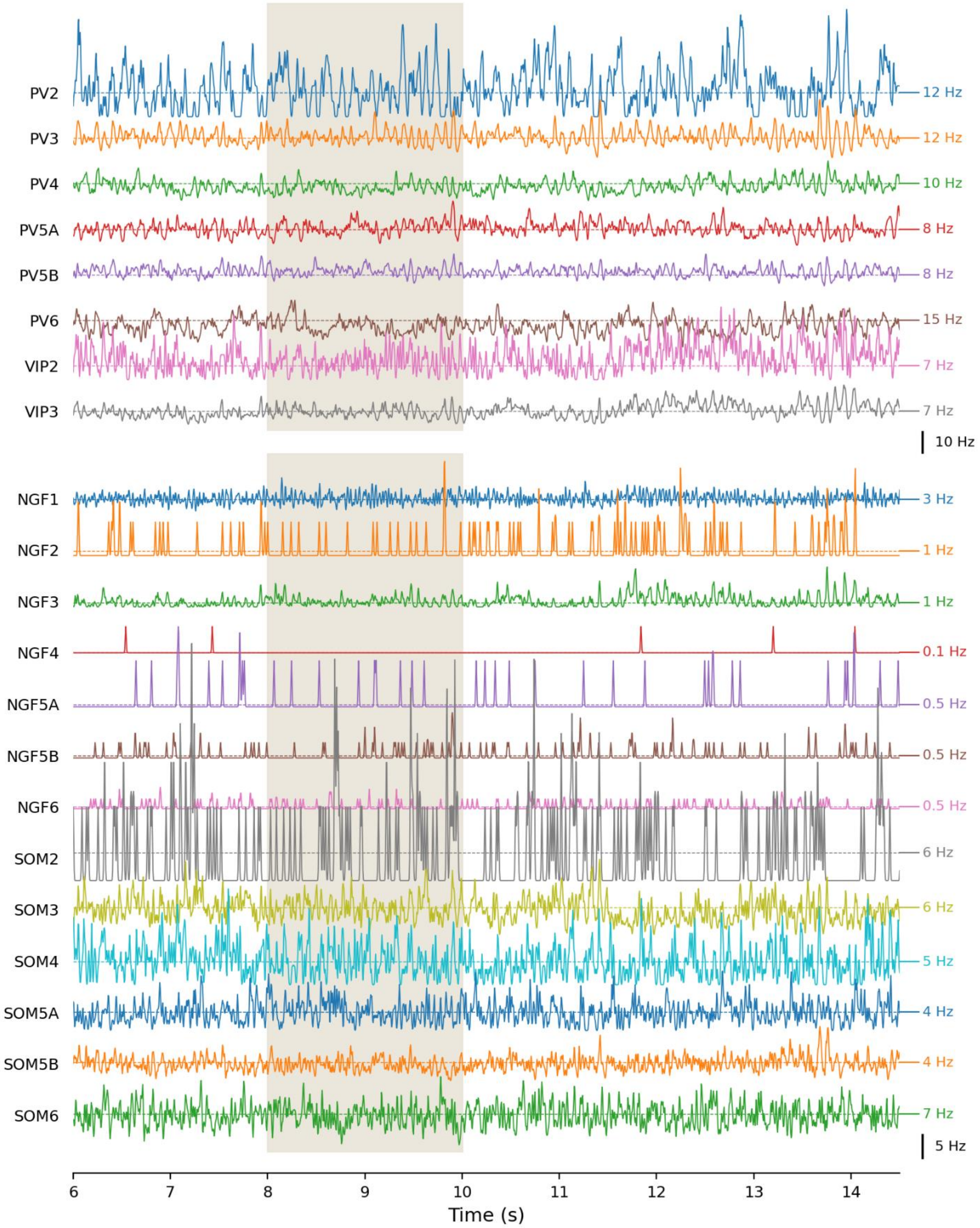


**Figure S4. Firing rate dynamics in the full thalamocortical model (inhibitory populations).** Legend is the same as in **S3**.

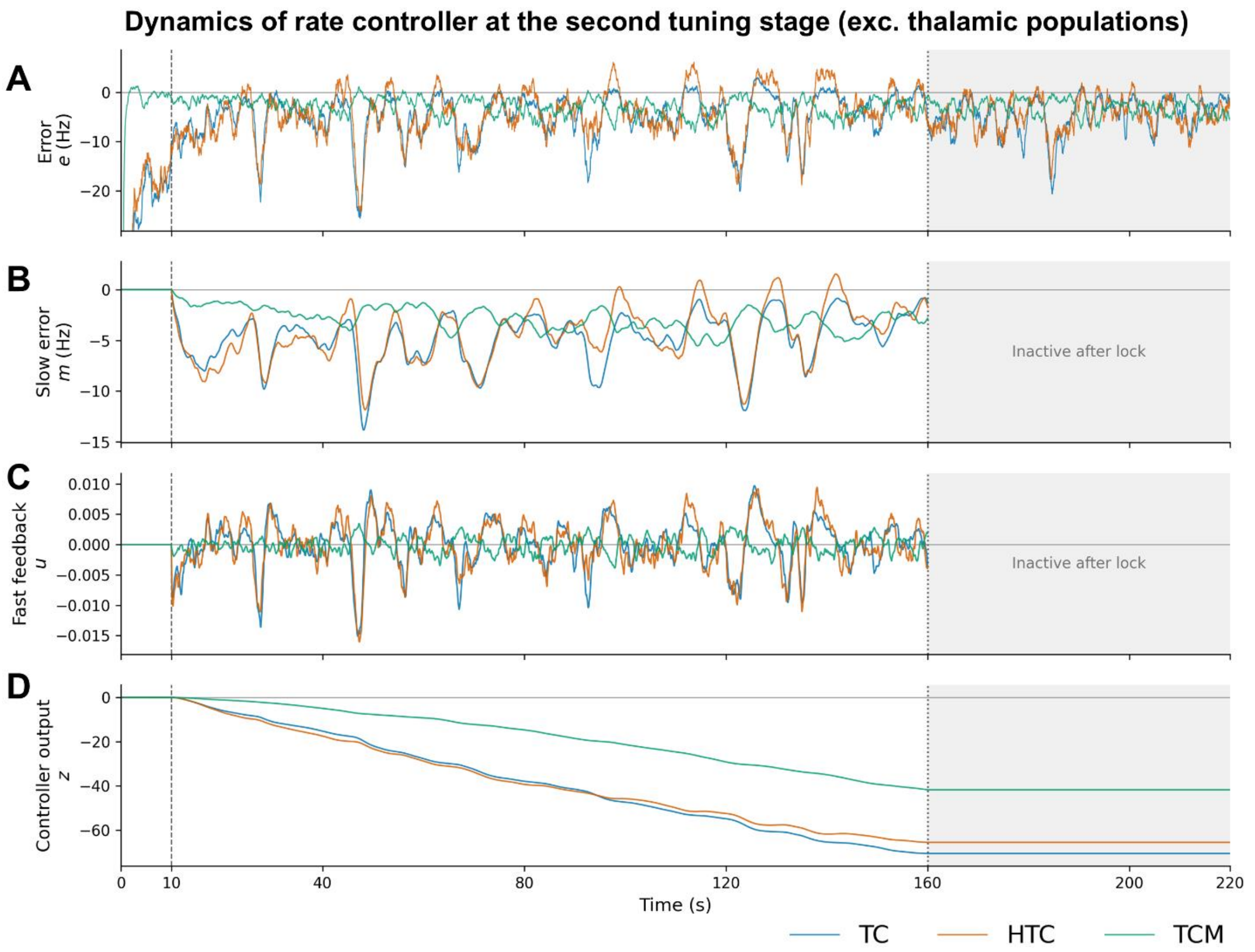


**Figure S5. Second-stage tuning in excitatory thalamic populations.** Traces show the time course of variables related to feedback firing rate controllers for the excitatory thalamic populations (blue – TC; orange – HTC; green – TCM) in one model realization. **(A)** Firing rate error: $e_p(t) = r_p^0 - r_p(t)$, where $r_p^0$ is the target rate and $r_p(t)$ is the filtered population rate. **(B)** Slow error component $m_p(t)$, computed by smoothing $e_p(t)$ with a 2.5-s exponential kernel. **(C)** Fast feedback component, $u_p(t)$, which responds to short-term error fluctuations. **(D)** Current-correction signal, $z_p$, converted to injected current as $I_p(t) = 0.005 \times z_p(t)$ [nA]. Feedback control begins at 10 s (dashed line). Correction $z_p$ is fixed at its learned value at 160 s (dotted line), for the remainder of the simulation (gray shading); variables $m_p$ and $u_p$ are no longer used after the locking. All samples were recorded every 25 ms and plotted without additional smoothing.

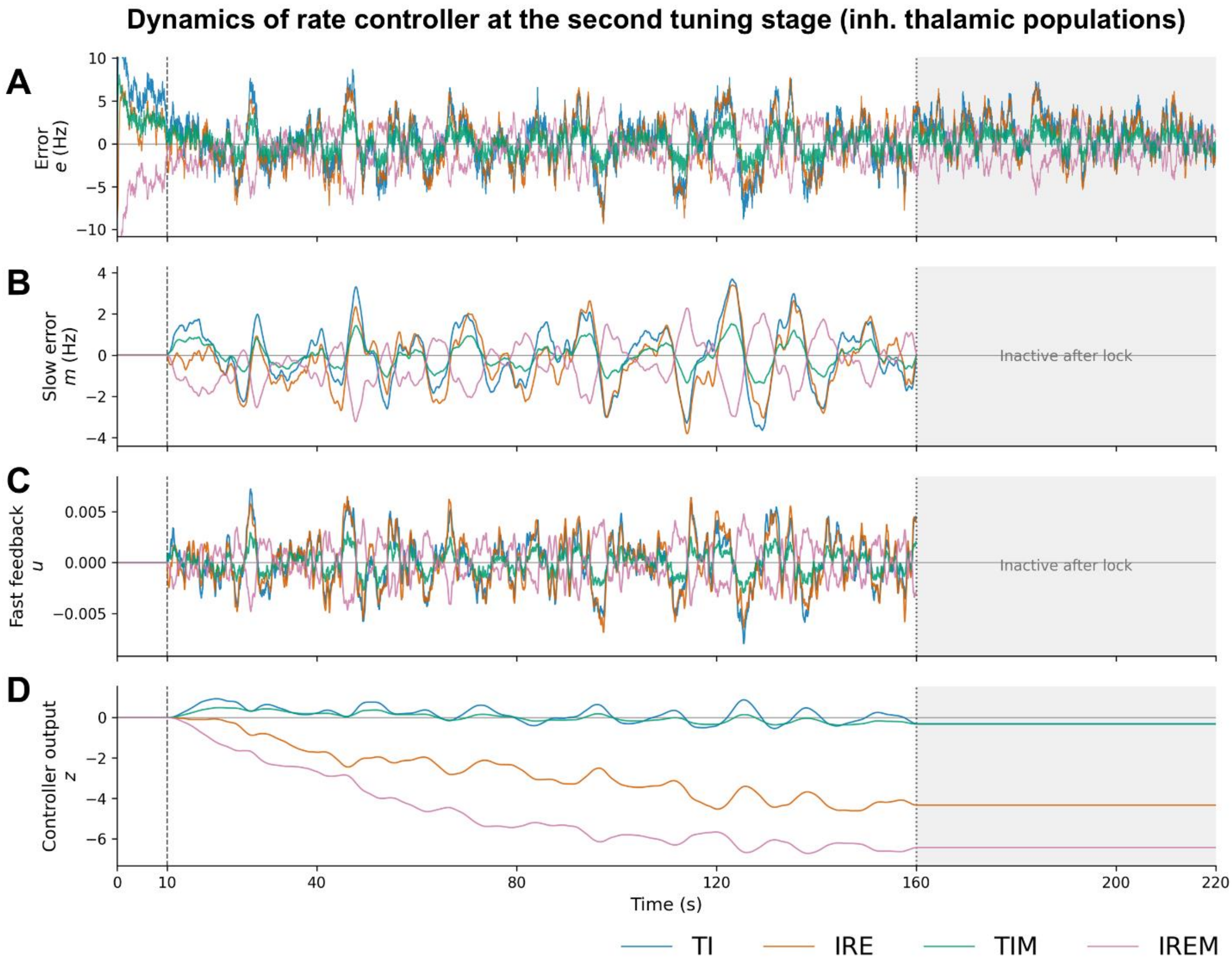


**Figure S6. Second-stage tuning in inhibitory thalamic populations.** Traces show the time course of variables related to feedback firing rate controllers for the inhibitory thalamic populations (blue – TI; orange – IRE; green – TIM, purple – IREM) in one model realization. The legend is the same as in **Fig. S5**.

### Tables

#### Table S1. Target firing rates.

| Population | Rate, Hz | Population | Rate, Hz |
|---|---|---|---|
| **NGF1** | 3 | **NGF5A** | 0.5 |
| **IT2** | 2 | **IT5B** | 5 |
| **PV2** | 12 | **CT5B** | 2 |
| **SOM2** | 6 | **PT5B** | 5 |
| **VIP2** | 7 | **PV5B** | 8 |
| **NGF2** | 1 | **SOM5B** | 4 |
| **IT3** | 2 | **VIP5B** | 0.2 |
| **PV3** | 12 | **NGF5B** | 0.5 |
| **SOM3** | 6 | **IT6** | 3 |
| **VIP3** | 7 | **CT6** | 2 |
| **NGF3** | 1 | **PV6** | 15 |
| **ITS4** | 2 | **SOM6** | 7 |
| **ITP4** | 2 | **VIP6** | 0.2 |
| **PV4** | 10 | **NGF6** | 0.5 |
| **SOM4** | 5 | **TC** | 8 |
| **VIP4** | 0.2 | **HTC** | 8 |
| **NGF4** | 0.1 | **TCM** | 2 |
| **IT5A** | 5 | **TI** | 15 |
| **CT5A** | 2 | **TIM** | 10 |
| **PV5A** | 8 | **IRE** | 15 |
| **SOM5A** | 4 | **IREM** | 10 |
| **VIP5A** | 0.2 | | |

#### Table S2. Tonic offset currents for setting the resting state.

| | Current, nA | | Current, nA |
|---|---|---|---|
| **IT2** | 0.183 | **IT6** | 0.096 |
| **IT3** | 0.207 | **CT6** | 0.281 |
| **ITS4** | 0.215 | **TC** | 4 |
| **ITP4** | 0.215 | **HTC** | 4 |
| **IT5A** | 0.099 | **TCM** | 6 |
| **CT5A** | 0.281 | **TI** | 0.03 |
| **IT5B** | 0.107 | **TIM** | 0.03 |
| **CT5B** | 0.281 | **IRE** | 0.2 |
| **PT5B** | 0.143 | **IREM** | 0.2 |

**Table S3. Background-input rates and synaptic weights.**

| | rxe [Hz] | rxi [Hz] | wxe [μS] | wxi [μS] | | rxe [Hz] | rxi [Hz] | wxe [μS] | wxi [μS] |
|---|---|---|---|---|---|---|---|---|---|
| **IT2** | 8776 | 939 | 0.31 | 1.17 | **SOM6** | 3419 | 1335 | 0.26 | 0.54 |
| **IT3** | 9575 | 1560 | 0.31 | 1.16 | **VIP2** | 2802 | 1024 | 0.31 | 1.03 |
| **ITP4** | 8385 | 1620 | 0.31 | 1.19 | **VIP3** | 2930 | 1054 | 0.31 | 1.03 |
| **ITS4** | 8400 | 1611 | 0.31 | 1.19 | **VIP4** | 1775 | 1599 | 0.31 | 1.03 |
| **IT5A** | 8705 | 2081 | 0.30 | 1.12 | **VIP5A** | 1625 | 622 | 0.31 | 1.03 |
| **IT5B** | 8763 | 2052 | 0.30 | 1.15 | **VIP5B** | 2117 | 672 | 0.31 | 1.03 |
| **IT6** | 7981 | 2006 | 0.29 | 1.09 | **VIP6** | 753 | 3407 | 0.31 | 1.03 |
| **CT5A** | 8287 | 1627 | 0.31 | 1.24 | **NGF1** | 6629 | 408 | 0.33 | 0.64 |
| **CT5B** | 8241 | 1636 | 0.31 | 1.24 | **NGF2** | 3451 | 708 | 0.33 | 0.64 |
| **CT6** | 7801 | 1632 | 0.31 | 1.24 | **NGF3** | 3790 | 765 | 0.33 | 0.64 |
| **PT5B** | 8553 | 1486 | 0.32 | 1.10 | **NGF4** | 2732 | 1090 | 0.33 | 0.64 |
| **PV2** | 5411 | 1578 | 0.25 | 1.99 | **NGF5A** | 3725 | 568 | 0.33 | 0.64 |
| **PV3** | 5405 | 1582 | 0.25 | 1.99 | **NGF5B** | 3649 | 532 | 0.33 | 0.64 |
| **PV4** | 5492 | 653 | 0.25 | 1.99 | **NGF6** | 3211 | 718 | 0.33 | 0.64 |
| **PV5A** | 5595 | 341 | 0.25 | 1.99 | **TC** | 8573 | 764 | 0.19 | 1.08 |
| **PV5B** | 5685 | 344 | 0.25 | 1.99 | **HTC** | 8130 | 954 | 0.19 | 1.08 |
| **PV6** | 5561 | 781 | 0.25 | 1.99 | **TCM** | 9361 | 650 | 0.19 | 1.08 |
| **SOM2** | 3066 | 3050 | 0.26 | 0.54 | **TI** | 8526 | 548 | 0.14 | 1.11 |
| **SOM3** | 3209 | 3091 | 0.26 | 0.54 | **TIM** | 8814 | 497 | 0.14 | 1.11 |
| **SOM4** | 3381 | 2583 | 0.26 | 0.54 | **IRE** | 8817 | 442 | 0.28 | 1.65 |
| **SOM5A** | 3268 | 1928 | 0.26 | 0.54 | **IREM** | 9709 | 390 | 0.28 | 1.65 |
| **SOM5B** | 3256 | 1955 | 0.26 | 0.54 | | | | | |

Excitatory and inhibitory Poisson input rates rxe, rxi were selected by open-loop tuning in the fully decoupled circuit. The corresponding synaptic weights wxe, wxi were fixed so that an isolated background spike produced a somatic voltage deflection of approximately 0.5 mV in absolute amplitude at rest.

**Table S4. Thalamic corrections obtained by secondary tuning.**

| | Median, nA | Mean, nA | Std., nA |
|---|---|---|---|
| **TC** | -0.2351 | -0.2541 | 0.1049 |
| **HTC** | -0.1580 | -0.1342 | 0.1440 |
| **TCM** | -0.2088 | -0.2217 | 0.0580 |
| **TI** | -0.0005 | -0.0005 | 0.0011 |
| **TIM** | -0.0009 | -0.0011 | 0.0010 |
| **IRE** | -0.0193 | -0.0200 | 0.0036 |
| **IREM** | -0.0308 | -0.0303 | 0.0082 |

**Table S5. Cortical firing rate errors across model configurations.**

| Population | Target rate (Hz) | Error, Hz | | | | |
|---|---|---|---|---|---|---|
| | | Isolated cortex | | | | Thalamocortical |
| | | Original | All-surr. | E-E surr. | E-E faded | E-E faded |
| **IT2** | 2 | -0.57 ± 0.15 | -0.12 ± 0.33 | -0.34 ± 0.14 | -0.38 ± 0.15 | -0.49 ± 0.18 |
| **IT3** | 2 | -0.02 ± 0.01 | -0.08 ± 0.12 | 0.03 ± 0.01 | 0.03 ± 0.01 | 0.01 ± 0.03 |
| **ITP4** | 2 | -0.04 ± 0.04 | -0.09 ± 0.09 | 0.00 ± 0.03 | -0.02 ± 0.04 | 0.05 ± 0.10 |
| **ITS4** | 2 | -0.03 ± 0.03 | -0.11 ± 0.10 | 0.00 ± 0.02 | 0.00 ± 0.02 | 0.05 ± 0.06 |
| **IT5A** | 5 | 1.91 ± 0.14 | 0.06 ± 0.14 | -0.13 ± 0.07 | -0.08 ± 0.06 | -0.10 ± 0.08 |
| **CT5A** | 2 | -0.08 ± 0.05 | 0.03 ± 0.07 | 0.09 ± 0.04 | 0.12 ± 0.05 | 0.09 ± 0.05 |
| **IT5B** | 5 | 2.86 ± 0.15 | -0.08 ± 0.12 | 0.10 ± 0.05 | 0.17 ± 0.06 | 0.20 ± 0.07 |
| **CT5B** | 2 | -0.25 ± 0.05 | -0.08 ± 0.07 | -0.04 ± 0.04 | -0.01 ± 0.04 | -0.04 ± 0.03 |
| **PT5B** | 5 | 0.36 ± 0.12 | 0.05 ± 0.17 | 0.33 ± 0.09 | 0.45 ± 0.11 | 0.36 ± 0.14 |
| **IT6** | 3 | -0.10 ± 0.03 | 0.02 ± 0.07 | 0.01 ± 0.03 | 0.00 ± 0.03 | 0.04 ± 0.07 |
| **CT6** | 2 | -0.08 ± 0.02 | -0.05 ± 0.04 | -0.03 ± 0.02 | -0.03 ± 0.02 | -0.01 ± 0.04 |
| **PV2** | 12 | 0.77 ± 1.81 | -0.57 ± 1.92 | 0.99 ± 1.61 | 1.02 ± 1.72 | 0.39 ± 1.92 |
| **PV3** | 12 | 0.61 ± 0.35 | -0.81 ± 1.52 | 0.63 ± 0.30 | 0.71 ± 0.37 | 0.27 ± 0.69 |
| **PV4** | 10 | 0.83 ± 0.58 | -0.84 ± 1.00 | -0.61 ± 0.42 | -0.74 ± 0.48 | 1.52 ± 3.21 |
| **PV5A** | 8 | 1.61 ± 0.72 | -0.96 ± 0.62 | 0.40 ± 0.61 | 0.43 ± 0.56 | 0.32 ± 0.92 |
| **PV5B** | 8 | 2.13 ± 0.50 | -0.14 ± 0.69 | 0.95 ± 0.38 | 1.06 ± 0.44 | 1.05 ± 0.64 |
| **PV6** | 15 | 1.21 ± 1.21 | -0.59 ± 1.22 | -1.27 ± 1.08 | -1.14 ± 1.08 | 1.82 ± 3.41 |
| **SOM2** | 6 | -2.35 ± 0.79 | -0.73 ± 0.99 | -0.68 ± 1.10 | -0.79 ± 1.20 | -1.18 ± 1.26 |
| **SOM3** | 6 | -2.10 ± 0.38 | -0.43 ± 0.51 | -0.31 ± 0.31 | -0.40 ± 0.45 | -0.89 ± 0.57 |
| **SOM4** | 5 | 0.68 ± 0.46 | -0.07 ± 0.48 | 0.17 ± 0.37 | 0.20 ± 0.44 | 0.88 ± 1.22 |
| **SOM5A** | 4 | 1.00 ± 0.39 | -0.39 ± 0.20 | -0.09 ± 0.29 | -0.02 ± 0.29 | -0.15 ± 0.31 |
| **SOM5B** | 4 | 0.85 ± 0.20 | -0.40 ± 0.20 | -0.20 ± 0.14 | -0.13 ± 0.14 | 0.31 ± 0.68 |
| **SOM6** | 7 | 0.63 ± 0.31 | -0.33 ± 0.24 | -0.13 ± 0.27 | -0.09 ± 0.26 | -0.20 ± 0.23 |
| **VIP2** | 7 | 10.32 ± 1.75 | -0.63 ± 0.85 | 0.95 ± 0.75 | 1.56 ± 0.77 | 3.10 ± 2.33 |
| **VIP3** | 7 | 10.20 ± 1.34 | -0.83 ± 0.79 | 0.81 ± 0.90 | 1.45 ± 1.19 | 2.68 ± 2.09 |
| **VIP4** | 0.2 | 0.13 ± 0.11 | -0.04 ± 0.10 | -0.04 ± 0.09 | -0.01 ± 0.09 | -0.08 ± 0.09 |
| **VIP5A** | 0.2 | -0.18 ± 0.03 | -0.18 ± 0.02 | -0.14 ± 0.06 | -0.13 ± 0.06 | -0.16 ± 0.03 |
| **VIP5B** | 0.2 | -0.06 ± 0.08 | -0.05 ± 0.06 | 0.02 ± 0.11 | 0.05 ± 0.14 | 0.00 ± 0.10 |
| **VIP6** | 0.2 | -0.17 ± 0.03 | -0.04 ± 0.21 | -0.14 ± 0.03 | -0.13 ± 0.04 | -0.11 ± 0.14 |
| **NGF1** | 3 | 0.20 ± 0.04 | 0.04 ± 0.09 | 0.11 ± 0.04 | 0.11 ± 0.05 | 0.17 ± 0.07 |
| **NGF2** | 1 | 0.62 ± 0.21 | -0.12 ± 0.23 | 0.08 ± 0.19 | 0.19 ± 0.21 | 0.30 ± 0.20 |
| **NGF3** | 1 | 0.73 ± 0.17 | 0.00 ± 0.11 | 0.19 ± 0.13 | 0.28 ± 0.18 | 0.41 ± 0.16 |
| **NGF4** | 0.1 | -0.05 ± 0.04 | -0.07 ± 0.03 | -0.06 ± 0.03 | -0.06 ± 0.03 | -0.03 ± 0.03 |
| **NGF5A** | 0.5 | 0.12 ± 0.14 | -0.04 ± 0.15 | 0.12 ± 0.11 | 0.12 ± 0.13 | 0.04 ± 0.18 |
| **NGF5B** | 0.5 | 0.05 ± 0.18 | -0.07 ± 0.13 | 0.08 ± 0.13 | 0.11 ± 0.14 | -0.01 ± 0.17 |
| **NGF6** | 0.5 | 0.02 ± 0.06 | -0.08 ± 0.04 | -0.07 ± 0.04 | -0.06 ± 0.05 | -0.08 ± 0.04 |

Each condition reports the mean signed error (population firing rate minus target) ± sample standard deviation across 15 random model realizations. Original - fully recurrent connectivity; All-surr. - all connections are surrogate; E-E surr. - excitatory-to-excitatory connections are surrogate, others are recurrent; E-E faded - excitatory-to-excitatory connections are faded from surrogate to recurrent.

**Table S6. Thalamic firing rate errors across model configurations.**

| Population | Target rate (Hz) | Error, Hz | | | | |
|---|---|---|---|---|---|---|
| | | Isolated thalamus | | | | Thalamocortical |
| | | Original | All-surr. | Corr. trial | Corr. median | Corr. median |
| **TC** | 8 | 9.09 ± 10.20 | 1.81 ± 2.91 | 1.26 ± 2.61 | 2.10 ± 7.22 | 7.65 ± 6.62 |
| **HTC** | 8 | 4.59 ± 6.67 | 0.29 ± 3.68 | 1.28 ± 1.82 | 0.73 ± 5.00 | 1.42 ± 7.16 |
| **TI** | 15 | -2.25 ± 3.47 | 0.37 ± 1.10 | 0.45 ± 1.46 | 0.53 ± 3.00 | -1.11 ± 2.45 |
| **IRE** | 15 | 0.58 ± 3.33 | -0.08 ± 1.32 | 0.48 ± 1.08 | 0.82 ± 3.11 | -0.82 ± 2.36 |
| **TCM** | 2 | 3.10 ± 1.85 | 0.53 ± 1.59 | 3.47 ± 0.89 | 3.92 ± 1.93 | 2.10 ± 1.44 |
| **TIM** | 10 | -1.30 ± 1.68 | 0.71 ± 1.63 | 0.27 ± 0.66 | 0.33 ± 1.46 | -0.35 ± 1.35 |
| **IREM** | 10 | 1.90 ± 2.91 | -0.02 ± 1.64 | -0.32 ± 0.88 | -0.37 ± 2.49 | 0.88 ± 2.07 |

Each condition reports the mean signed error (population firing rate minus target) ± sample standard deviation across 15 random model realizations, in Hz. Original – recurrent thalamic connectivity, no additional tonic corrections; All-surr. – all connections are surrogate; Corr. trial – recurrent thalamic connectivity with tonic corrections optimized separately for each model realization; Corr. median – recurrent thalamic connectivity with the median correction across realizations applied to each population.

**Table S7. Population-specific regions for background-rate search.**

| **Pop.** | $r_{xe}^{p,max}$ [kHz] | $r_{xi}^{p,max}$ [kHz] | $r_{xe}^{pc}$ [kHz] | $r_{xi}^{pc}$ [kHz] | $\Delta\, r_{xe}^{p}$ [kHz] | $\Delta\, r_{xi}^{p}$ [kHz] | $s_p$ | $r_p^{high}$ [Hz] | $\alpha_p$ | $\beta_p$ |
|---|---|---|---|---|---|---|---|---|---|---|
| **CT5A** | 25 | 5 | 2.75 | 0 | 6.35 | 4.05 | 0.178 | 10 | 0.3 | 0.1 |
| **CT5B** | 25 | 5 | 2.72 | 0 | 6.38 | 4.07 | 0.179 | 10 | 0.3 | 0.1 |
| **CT6** | 25 | 5 | 2.36 | 0 | 6.47 | 4.14 | 0.179 | 10 | 0.3 | 0.1 |
| **IT2** | 20 | 4 | 9.48 | 0 | 9.25 | 2.01 | 0.179 | 30 | 0.3 | 0.1 |
| **IT3** | 25 | 5 | 3.53 | 0 | 5.2 | 3.88 | 0.178 | 10 | 0.3 | 0.1 |
| **ITP4** | 25 | 5 | 2.72 | 0 | 6.13 | 4.08 | 0.180 | 10 | 0.3 | 0.1 |
| **ITS4** | 25 | 5 | 2.75 | 0 | 6.17 | 4.06 | 0.179 | 10 | 0.3 | 0.1 |
| **IT5A** | 25 | 5 | 1.73 | 0 | 6.1 | 4.99 | 0.218 | 10 | 0.3 | 0.1 |
| **IT5B** | 25 | 5 | 1.72 | 0 | 6.73 | 4.98 | 0.209 | 10 | 0.3 | 0.1 |
| **IT6** | 25 | 5 | 2.17 | 0 | 5.98 | 4.99 | 0.224 | 10 | 0.3 | 0.1 |
| **NGF1** | 15 | 3 | 10.37 | 0 | 12.32 | 0.82 | 0.148 | 10 | 0.3 | 0.1 |
| **NGF2** | 15 | 3 | 7.06 | 0 | 12.25 | 1.33 | 0.151 | 10 | 0.3 | 0.1 |
| **NGF3** | 15 | 3 | 7.12 | 0 | 11.83 | 1.36 | 0.156 | 10 | 0.3 | 0.1 |
| **NGF4** | 15 | 3 | 5.54 | 0 | 13.08 | 1.56 | 0.149 | 10 | 0.3 | 0.1 |
| **NGF5A** | 15 | 3 | 9.03 | 0 | 11.19 | 1.13 | 0.166 | 10 | 0.3 | 0.1 |
| **NGF5B** | 15 | 3 | 9.16 | 0 | 11.78 | 1.07 | 0.159 | 10 | 0.3 | 0.1 |
| **NGF6** | 15 | 3 | 7.16 | 0 | 11.69 | 1.36 | 0.157 | 10 | 0.3 | 0.1 |
| **PT5B** | 25 | 4 | 2.26 | 0 | 9.07 | 3.36 | 0.151 | 10 | 0.3 | 0.1 |
| **PV2** | 15 | 2.5 | 0 | 1.01 | 12.52 | 0.89 | 0.056 | 60 | 0.3 | 0.1 |
| **PV3** | 15 | 2.5 | 0 | 1.01 | 11.92 | 0.89 | 0.056 | 60 | 0.3 | 0.1 |
| **PV4** | 15 | 2.5 | 0 | 0.14 | 9.87 | 0.88 | 0.056 | 60 | 0.3 | 0.1 |
| **PV5A** | 15 | 2.5 | 2.76 | 0 | 8.52 | 0.73 | 0.057 | 60 | 0.3 | 0.1 |
| **PV5B** | 15 | 2.5 | 2.84 | 0 | 8.63 | 0.72 | 0.057 | 60 | 0.3 | 0.1 |
| **PV6** | 15 | 2.5 | 0 | 0.23 | 10.01 | 0.93 | 0.059 | 60 | 0.3 | 0.1 |
| **SOM2** | 10 | 5 | 0 | 1.03 | 6.25 | 3.14 | 0.300 | 60 | 0.3 | 0.1 |
| **SOM3** | 10 | 5 | 0 | 1.04 | 6.1 | 3.14 | 0.300 | 60 | 0.3 | 0.1 |
| **SOM4** | 10 | 5 | 0 | 0.48 | 5.86 | 3.17 | 0.303 | 60 | 0.3 | 0.1 |
| **SOM5A** | 10 | 5 | 0.48 | 0 | 5.52 | 3.07 | 0.309 | 60 | 0.3 | 0.1 |
| **SOM5B** | 10 | 5 | 0.4 | 0 | 5.45 | 3.1 | 0.310 | 60 | 0.3 | 0.1 |
| **SOM6** | 10 | 5 | 1.91 | 0 | 5.18 | 2.58 | 0.305 | 60 | 0.3 | 0.1 |
| **TC** | 15 | 3 | 8.11 | 0 | 14.53 | 1.6 | 0.120 | 8 | 0.8 | 0.8 |
| **HTC** | 15 | 3 | 9.3 | 0 | 16.84 | 1.68 | 0.128 | 8 | 0.8 | 0.8 |
| **TCM** | 15 | 3 | 7.68 | 0 | 9.24 | 1.25 | 0.110 | 2 | 0.8 | 0.8 |
| **TI** | 15 | 1.5 | 6.73 | 0 | 8.34 | 1.09 | 0.090 | 15 | 0.5 | 0.7 |
| **TIM** | 15 | 1.5 | 6.91 | 0 | 6.99 | 1.02 | 0.090 | 10 | 0.5 | 0.7 |
| **IRE** | 15 | 1.5 | 6.58 | 0 | 7.55 | 0.88 | 0.074 | 15 | 0.5 | 0.7 |
| **IREM** | 15 | 1.5 | 7.8 | 0 | 6.74 | 0.76 | 0.074 | 10 | 0.5 | 0.7 |
| **VIP2** | 10 | 5 | 1.18 | 0 | 9.78 | 1.51 | 0.158 | 60 | 0.3 | 0.1 |
| **VIP3** | 10 | 5 | 1.18 | 0 | 9.19 | 1.52 | 0.160 | 60 | 0.3 | 0.1 |
| **VIP4** | 10 | 5 | 0 | 0.12 | 9.64 | 1.84 | 0.172 | 60 | 0.3 | 0.1 |
| **VIP5A** | 10 | 5 | 3.61 | 0 | 10 | 0.99 | 0.139 | 60 | 0.3 | 0.1 |
| **VIP5B** | 10 | 5 | 3.64 | 0 | 9.35 | 0.99 | 0.141 | 60 | 0.3 | 0.1 |
| **VIP6** | 10 | 5 | 0 | 1.26 | 10.66 | 2.31 | 0.214 | 60 | 0.3 | 0.1 |

For each population $P$, the coarse scan sampled excitatory background input rates from 1 Hz to $r_{xe}^{p,max}$ and inhibitory background input rates from 1 Hz to $r_{xi}^{p,max}$. The fine-search parallelogram $Q_p$ is specified by its lower-right corner $(r_{xe}^{pc}, r_{xi}^{pc})$, horizontal width $\Delta\, r_{xe}^{p}$, vertical height $\Delta\, r_{xi}^{p}$, and slope $s_p$ of its slanted edges. The low- and high-rate contours used to construct $Q_p$ were extracted from the coarse firing rate map $R_p^b$ at population firing rates $r_p^{low}$ and $r_p^{high}$, respectively; $r_p^{low} = 0.5\ Hz$ for all populations. The width-expansion fraction $\alpha_p$ determines the horizontal width of $Q_p$. The right-edge shift fraction $\beta_p$ specifies a rightward displacement of the region.

**Table S8. Current cellular and inhibitory synaptic parameters that differ from Dura-Bernal et al. (2023a)**

| Model element | Population or pathway | Parameter or mechanism | Current setting |
|---|---|---|---|
| Cortical cell | IT2; IT3 | kdr.gbar, all sections | 1.5 × source-template value |
| Cortical cell | ITP4; IT5A; CT5A; IT5B; CT5B; IT6; CT6 | kdr.gbar, all sections | 3 × source-template value |
| Cortical cell | ITS4 | Sections | soma; Bdend; Adend1; Adend2; Adend3 |
| Thalamic cell | TC | cal mechanism | none |
| Thalamic cell | TC | htc.gmax | 1e-6 S/cm² |
| Thalamic cell | TC | ittc.gmax | 0.002 S/cm² |
| Thalamic cell | HTC | cal.gcalbar | 0.001 S/cm² |
| Thalamic cell | HTC | htc.gmax | 1e-6 S/cm² |
| Thalamic cell | HTC | ittc.gmax | 0.002 S/cm² |
| Inhibitory synapse | GABABCtx; GABABThal | tau1 | 41 ms |
| Inhibitory synapse | GABABCtx; GABABThal | tau2 | 642 ms |
| Inhibitory synapse | GABABCtx; GABABThal | Reversal potential | −105 mV |
| Inhibitory projection | SOM→cortical E | GABAASlow; GABABCtx | 0.9; 0.2 |
| Inhibitory projection | SOM→cortical I | GABAASlow | 1.0 |
| Inhibitory projection | NGF→cortical E | GABAA; GABABCtx | 0.5; 1.0 |
| Inhibitory projection | NGF→cortical I | GABAA | 1.0 |
| Inhibitory projection | Thalamic I→thalamic E | GABAASlow; GABABThal | 0.9; 0.4 |
| Inhibitory projection | Thalamic I→thalamic I | GABAASlow | 1.0; no GABAB component |
| Inhibitory projection | Thalamic I→cortical E | GABAASlow; GABABThal | 1.0; 0 |
| Inhibitory projection | Thalamic I→cortical I | GABAASlow | 1.0 |

Cortical E denotes IT2, IT3, ITP4, ITS4, IT5A, CT5A, IT5B, CT5B, PT5B, IT6, and CT6; cortical I denotes the PV, SOM, VIP, and NGF populations; thalamic E denotes TC, TCM, and HTC; and thalamic I denotes TI, TIM, IRE, and IREM. Values in the "Inhibitory projection" rows are unitless multipliers for the listed receptor components.

**Table S9. Pathway-specific postsynaptic section assignments**

| Presynaptic populations | Postsynaptic populations | Sections |
|---|---|---|
| Cortical E, SOM, or VIP | Cortical E | Adend1; Adend2; Adend3; Bdend |
| PV | Cortical E | Adend1; Bdend; soma |
| NGF1 | Cortical E | Adend3 |
| NGF2, NGF3, or NGF4 | Cortical E2–4 | Adend1; Adend2 |
| NGF2, NGF3, or NGF4 | Cortical E5–6 | Adend2 |
| NGF5A, NGF5B, or NGF6 | Cortical E5–6 | Adend1 |
| NGF5A, NGF5B, or NGF6 | Cortical E2–4 | Bdend |
| TC or HTC | Cortical E | Adend1; Bdend |
| TCM | Cortical E | Adend1; Adend2; Adend3 |
| Cortical E | PV, SOM, or NGF | dend |
| Cortical E | VIP | ori1; ori2; rad1; rad2 |
| NGF, PV, or SOM | PV | dend |
| VIP | PV | soma |
| PV or VIP | SOM | soma; dend |
| NGF or SOM | SOM | dend |
| NGF, PV, SOM, or VIP | NGF | dend |
| PV | VIP | soma; ori1; ori2; rad1 |
| SOM | VIP | ori1; ori2; rad1; rad2 |
| VIP | VIP | soma; ori1; ori2; rad1; rad2 |
| NGF | VIP | ori1; ori2; rad1; rad2 |
| Cortical E or any thalamic/reticular population | Any thalamic/reticular population | soma |
| Any thalamic/reticular population | PV, SOM, or NGF | dend |
| Any thalamic/reticular population | VIP | ori1; ori2; rad1; rad2 |

Cortical E2–4 denotes IT2, IT3, ITP4, and ITS4. Cortical E5–6 denotes IT5A, CT5A, IT5B, CT5B, PT5B, IT6, and CT6. Any thalamic/reticular population denotes TC, TCM, HTC, TI, TIM, IRE, IREM. PV, SOM, VIP, and NGF denote all layer-specific populations of the corresponding inhibitory class. Section names: Adend1, Adend2, and Adend3 – successive apical dendritic sections (proximal to distal); Bdend – basal dendrite; dend – single dendritic section of the PV, SOM, and NGF cells; ori1, ori2, rad1, rad2 – dendritic sections of VIP cells.

**Table S10. Parameters of the second tuning stage**

| | | |
|---|---|---|
| Time-unit conversion | $c_t$ | 1000 ms s$^{-1}$ |
| Rate-estimator time constant | $\tau_r$ | 100 ms |
| Slow-error time constant | $\tau_m$ | 2500 ms |
| Fast-branch time constant | $\tau_u$ | 100 ms |
| Slow integrator gain | $k_s$ | 0.0001 ms$^{-1}$ |
| Fast-branch gain | $k_u$ | 0.001 |
| Error dead-zone half-width | $\varepsilon$ | 0.1 Hz |
| Feedback start | $t_0$ | 10 s |
| Locking time | $t_{\mathrm{lock}}$ | 160 s |
| Current gain | $g_\mu$ | 0.005 nA Hz$^{-1}$ |
